# Structural basis of canonical TIR-NLR activation in plant innate immunity

**DOI:** 10.64898/2026.02.03.703613

**Authors:** Natsumi Maruta, Weixi Gu, Bryan Ying Jie Lim, Mitchell Sorbello, Dalton Heng Yong Ngu, Chacko Jobichen, Jeffrey D. Nanson, Yan Li, Jian Chen, Megan A. Outram, Maud Bernoux, Motiur Rahman, Thi Danh Vu, Hongyi Xu, Lei Wang, Kayden Kwah, Hayden Burdett, Mehdi Mobli, Thomas Ve, Jeffrey G. Ellis, Peter A. Anderson, Simon J. Williams, Peter N. Dodds, Bostjan Kobe

**Author notes:** Correspondence (BK), (NM).

## Abstract

In plants, intracellular NLRs (nucleotide-binding leucine-rich repeat receptors) detect pathogen effector proteins, form oligomeric resistosomes, and activate ETI (effector-triggered immunity). NLRs contain N-terminal signaling, central NB-ARC (nucleotide-binding) and C-terminal LRR (leucine-rich repeat) domains. NLRs with N-terminal TIR (Toll/interleukin-1 receptor) domains (TNLs) hydrolyze NAD^+^ (nicotinamide adenine dinucleotide) to generate signaling molecules. We determined cryo-EM structures of flax M, a canonical non C-JID (C-terminal jellyroll/Ig-like domain) TNL, in both monomeric autoinhibited conformation, and tetrameric resistosome after activation by its rust fungal effector AvrM-A. AvrM-A homodimers dissociate into monomers to bind directly to the LRR and NB-ARC domains in the M resistosome. The resistosome structure includes a non-hydrolyzable NAD^+^ analogue, revealing the substrate NAD^+^ recognition mechanism by the TIR domains. M cleaves NAD^+^ and generates the same signaling compounds as the related flax TNL, L6. Our findings explain the mechanism of TNL signaling, and provide a basis for rational engineering of disease-resistant crops.

## Introduction

Plant diseases caused by pathogenic microbes pose a threat to food security, as they lead to severe yield losses in major crops. To combat pathogens, plants have evolved two types of innate immune receptors. PRRs (pattern recognition receptors) on the plasma membrane detect PAMPs (pathogen-associated molecular patterns) and activate PTI (PAMP-triggered immunity) to a broad spectrum of non-adapted pathogens. By contrast, intracellular NLRs (nucleotide-binding leucine-repeat receptors) recognize specific effector molecules secreted by pathogens and activate ETI (effector-triggered immunity), featuring localized programmed cell death as a hallmark of HR (hypersensitive response) ^1,2^. NLRs are found in plants and animals and share a conserved tripartite domain architecture: (i) a C-terminal LRR (leucine-rich repeat) domain involved in effector recognition, (ii) a central nucleotide-binding and oligomerization domain (termed NB-ARC in plant NLRs and NACHT in animal NLRs), serving as a molecular switch, with bound ADP (adenosine diphosphate) and ATP (adenosine triphosphate) representing the off and on state, respectively; and (iii) an N-terminal domain involved in signaling. Based on the N-terminal TIR (Toll/interleukin-1 receptor) and CC (coiled-coil) domains, plant NLRs are classified as TNLs (TIR-NLRs) and CNLs (CC-NLRs), respectively. Upon effector recognition, plant NLRs form oligomeric complexes termed resistosomes and activate downstream innate immune signaling pathways. Some CNLs can function alone, while TNLs require downstream helper NLRs known as RNLs (resistance to powdery mildew 8-like NLRs; CC_R_-NLRs). Additionally, in Asterids, a distinct clade of CNLs known as NRCs (NLRs required for cell death; helper NLRs) functions downstream of multiple CNLs, forming an NLR network ^3^.

Recent cryo-EM (cryogenic electron microscopy) structural studies reveal that CNLs and NRCs are monomeric and dimeric in their inactive states, respectively ^4–6^. Effector recognition induces CNLs, NRCs and CC_G10_-type CNLs to form pentameric, hexameric and octameric resistosomes, respectively ^7–12^. These oligomeric complexes function as membrane-associated calcium influx channels and trigger HR ^7,8,10,13^. On the other hand, the TNLs RPP1 from *Arabidopsis thaliana* and ROQ1 from *Nicotiana benthamiana* form tetrameric resistosomes, with the assembled TIR domains functioning as enzymes that hydrolyze NAD^+^ (nicotinamide adenine dinucleotide) ^14–17^. RPP1 and ROQ1 belong to a subclass of TNLs in flowering plants that additionally carry a C-JID (C-terminal jellyroll/Ig-like domain), which is found in approximately 50% of TNLs from dicot genomes and is essential for effector recognition ^15,16^. TNL-mediated NAD^+^ hydrolysis produces various signaling molecules that bind to the EDS1 (enhanced disease susceptibility 1) family of proteins, which in turn recruit RNLs, such as NRG1 (N requirement gene 1) and ADR1 (activated disease resistance 1), forming distinct heterotrimers ^18–23^. NRG1 and ADR1 oligomerize to form calcium influx channels, ultimately leading to elicitation of host cell death and disease resistance ^24^. Given the diversity of plant TNLs, it remains to be examined if tetramer formation is a conserved mechanism. Furthermore, structures of canonical, non-C-JID type, TNLs in their resting and activated states are unknown.

Rust fungi represent the largest group of phytopathogens causing devastating disease epidemics in agricultural crops ^25^. The flax rust fungus (*Melampsora lini*) infects cultivated flax (*Linum usitatissimum*), which is grown for linen fibre and oil production ^26^. Historically, the genetic basis of the flax-rust interaction has been extensively studied, founding the concept of gene-for-gene resistance underlying ETI ^27^. Several flax TNL-encoding genes, and their matching rust fungal effectors, have been identified and characterized ^28,29^, including the *M* resistance gene ^30^ and the corresponding AvrM effector ^31^. Therefore, this system serves as a fundamental model to study the molecular mechanisms of ETI. The flax M protein has a prototypical domain architecture, consisting of the TIR, NB-ARC (comprising of three subdomains, NBD (nucleotide-binding domain), HD1 (helical domain 1), and WHD (winged-helix domain)), and LRR domains ^30^ and directly interacts with the AvrM-A effector to trigger resistance ^32^. It is unclear how M maintains its resting state in the absence of the effector and how interaction with AvrM-A leads to M activation. Whether M possesses enzymatic activity and what associated signaling products are generated also remains unknown.

In this study, we expressed and purified a recombinant M protein that was predominantly in a monomeric state. Upon incubation with AvrM-A, ATP and an NAD^+^ analogue, M assembled into a tetramer, resembling the RPP1 and ROQ1 resistosomes. We determined the cryo-EM structure of the M resistosome at 2.7 Å resolution, revealing details of effector interaction, the nucleotide-binding state of the NB-ARC domain, as well as the TIR domain substrate binding site. Upon incubation with its effector, ATP and NAD^+^, M exhibited NADase activity and generated signaling compounds in an equivalent manner to a closely related flax TNL, L6. Our cryo-EM 3D reconstruction of the monomeric M protein revealed its closed conformation that resembles the autoinhibited state of CNLs, such as ZAR1 and NRC2, suggesting the conservation of the inactive conformation of plant NLRs. Overall, our study defines the structural and molecular basis of canonical TNL function.

## Results

### Purification of monomeric M protein and effector-induced resistosome formation *in vitro*

Near-full-length M protein (residues 21-1305, lacking the N-terminal signal anchor sequence) ^33^, with an N-terminal 6xHis-Twin-Strep-tag, was expressed in *Pichia pastoris* cells and purified by tandem-affinity purification, followed by SEC (size exclusion chromatography). The fractions collected from the largest peak in the SEC chromatogram (80-84 mL elution volume, Figure 1A) contained protein of the expected molecular weight of 150 kDa, with high purity, as determined by SDS-PAGE (sodium dodecyl sulfate–polyacrylamide gel electrophoresis; Figure 1B) and were used for further analysis. NanoDSF (nano differential scanning fluorimetry) measurements showed that the M protein melting temperature (T_m_, at which 50% of protein is unfolded), was at 56.8 °C (Figure S1), suggesting the protein is stable at room temperature. Mass photometry analysis showed that the protein size in solution was approximately 150 kDa (Figure 1C), corresponding to the predicted size of its monomeric form. Homogeneous sample distribution of the monomeric M protein was confirmed by negative-stain EM and cryo-EM analysis (Figure 1D).

**Figure 1.**
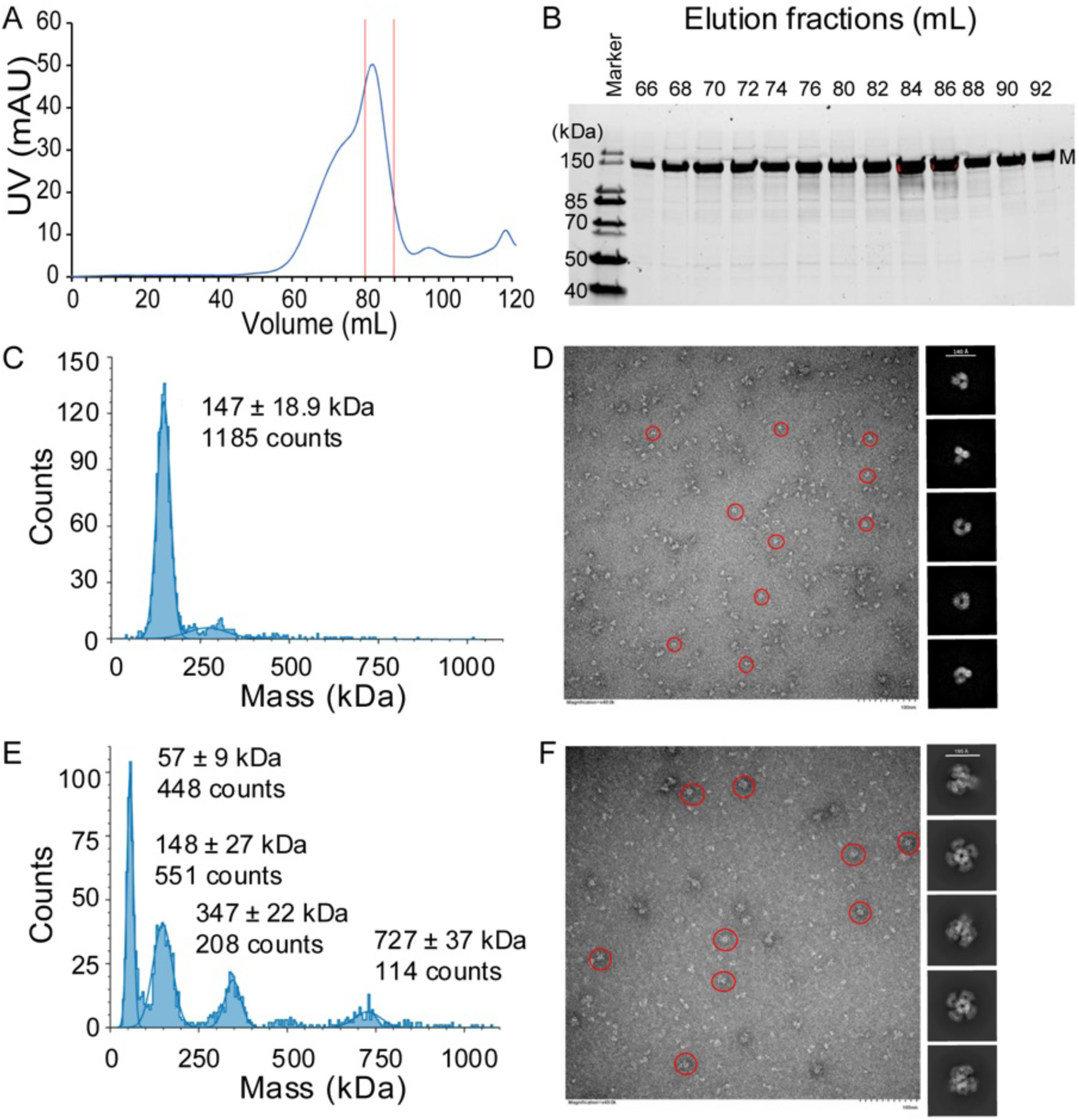
Purification and biophysical characterization of the flax M protein. Recombinant M protein was expressed in *Pichia pastoris* and purified using tandem affinity chromatography (StrepTactin XT and Ni-NTA resins), followed by SEC (size-exclusion chromatography). **(A)** SEC profile of full-length M protein on a HiLoad 16/600 Superose 6 pg (Cytiva) column. **(B)** Purity of the eluted fractions (66 - 92 mL) from the SEC column was determined by SDS-PAGE. Fractions from elution volumes 80 - 84 mL were used for further analysis. The Sypro Ruby stain was used to visualize the gel. **(C)** Mass photometry analysis; and **(D)** negative-stain EM (left panel) and cryo-EM 2D classification (right panel) of the monomeric M protein. **(E)** Mass photometry analysis of the M protein upon incubation with the avirulence effector AvrM-A and ATP (no NAD^+^ was added). **(F)** Negative-stain EM (left panel) and cryo-EM 2D classification (right panel) of the M protein, upon incubation with AvrM-A, ATP and 1AD. Negative-stain EM images for the protein samples were taken at 60,000x magnification. Scale bar = 100 nm. Representative particles of the monomer and M resistosome are enclosed in red circles.

The studies that led to the available structures of TNL resistosomes employed co-expression of TNLs and cognate effectors in plants or insect cells ^15,16^. In this study, we reconstituted the M resistosome *in vitro*, by mixing purified M and AvrM-A proteins. Mass photometry analysis showed that incubation of M with AvrM-A and ATP, in the presence or absence of NAD^+^ or a non-hydrolyzable NAD^+^ analogue (1AD, 5-iodo-isoquinoline ^34^), led to mixture of protein populations with different sizes, including ∼60 kDa, ∼150 kDa, ∼350 kDa and ∼730 kDa, which presumably corresponded to AvrM-A dimers, M monomers, M dimers and M tetramers, respectively (Figure 1E; Figure S2). The 730 kDa protein complex approximately matched the size of four molecules of M (150 kDa) and AvrM-A (28 kDa) in a 1:1 stoichiometric ratio, and was most abundant in the presence of 1AD (Figure S2). The M resistosome was observed under negative-stain EM and cryo-EM (Figure 1F), revealing the clover leaf-like structure resembling those of RPP1 and ROQ1 resistosomes. Therefore, we prepared the M resistosome in the presence of 1AD for cryo-EM single particle analysis (Figure S3).

### Cryo-EM structure of the M resistosome

M is a canonical TNL (without a C-JID) and contains 28 LRR motifs (Figure 2, 3; Figure S4), the longest among all the reported NLR structures. A distinct feature of M^LRR^ is that, in the C-terminal LRR domain, there are two copies of a tandem repeat (TR) sequence, each comprising five LRR motifs, with TR1 spanning LRR17 to LRR21 and TR2 spanning LRR22 to LRR26 ^30^ (Figure 3A, Figure S4). The cognate effector, AvrM-A, consists of coiled-coil domains comprising 11 α helices (PDB (Protein Data Bank): 4BJN ^35^; Figure S5A). Each M protomer is bound to a single AvrM-A molecule (Figure 2B, C). The tetramer formation is mediated by the NB-ARC domain, allowing the TIR domains to assemble into a dimer of dimers, stabilized by the “AE and BE” interfaces (see below) ^15,16,34,36,37^ (Figure 2B, C).

**Figure 2.**
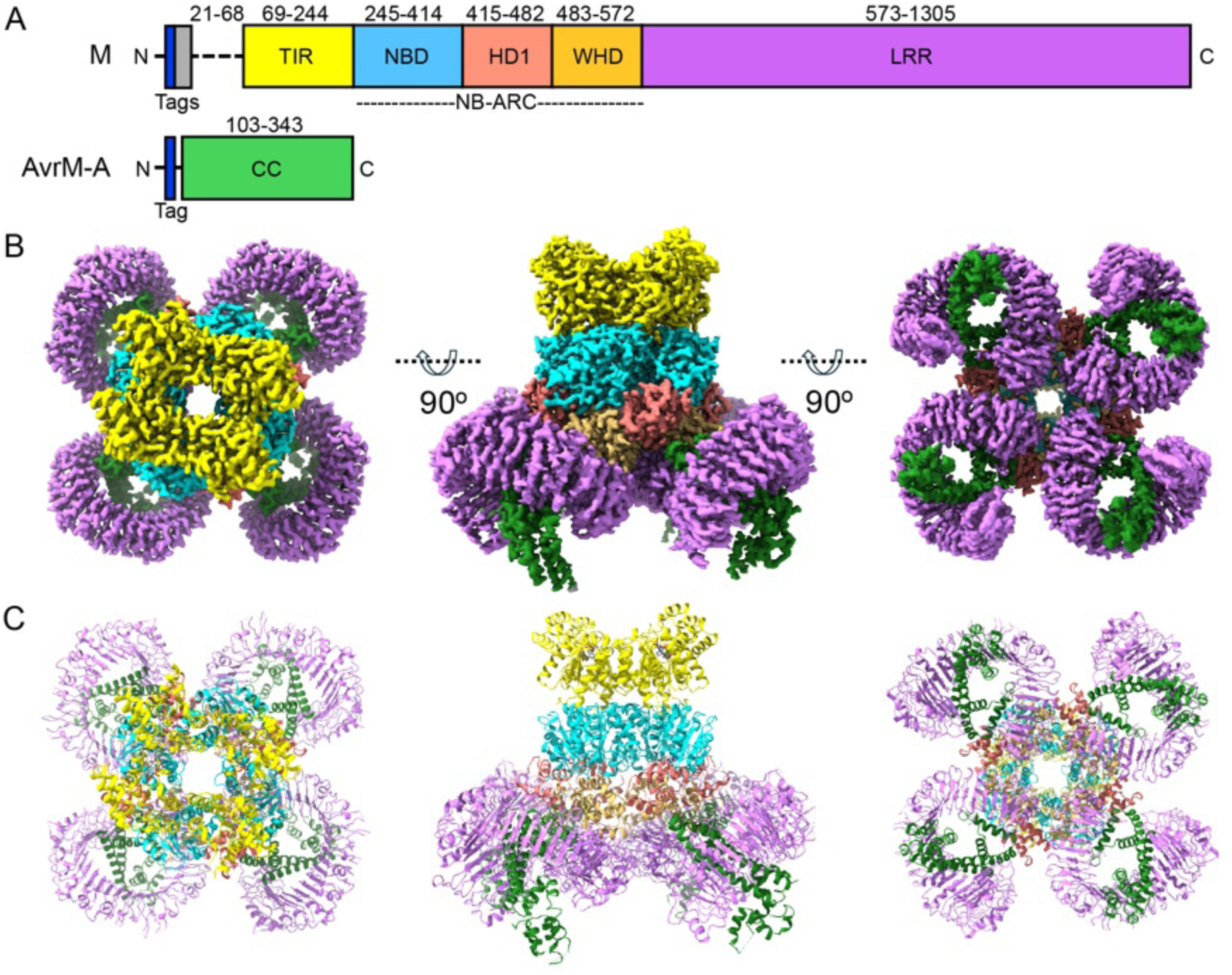
Cryo-EM structure of the M resistosome. **(A)** Schematic representation of M (top) and the effector AvrM-A (bottom) constructs. Tags for recombinant protein purification include a Twin-Strep tag (grey) and/or a 6xHis tag (blue). Residue numbers corresponding to the different structural domains are labeled. Residues 1-20 from the M protein and residues 1-102 from the AvrM-A protein have been excluded, as they correspond to the signal sequences. M (residues 21-68) corresponds to a disordered region. **(B)** Cryo-EM composite density map of the tetrameric M resistosome bound to AvrM-A. **(C)** The atomic model of the M resistosome. The M:AvrM-A resistosome complex is colored based on the domains, as described in **(A)**. Three different views of the protein complex are shown: “top” (left), “side” (middle) and “bottom” (right).

**Figure 3.**
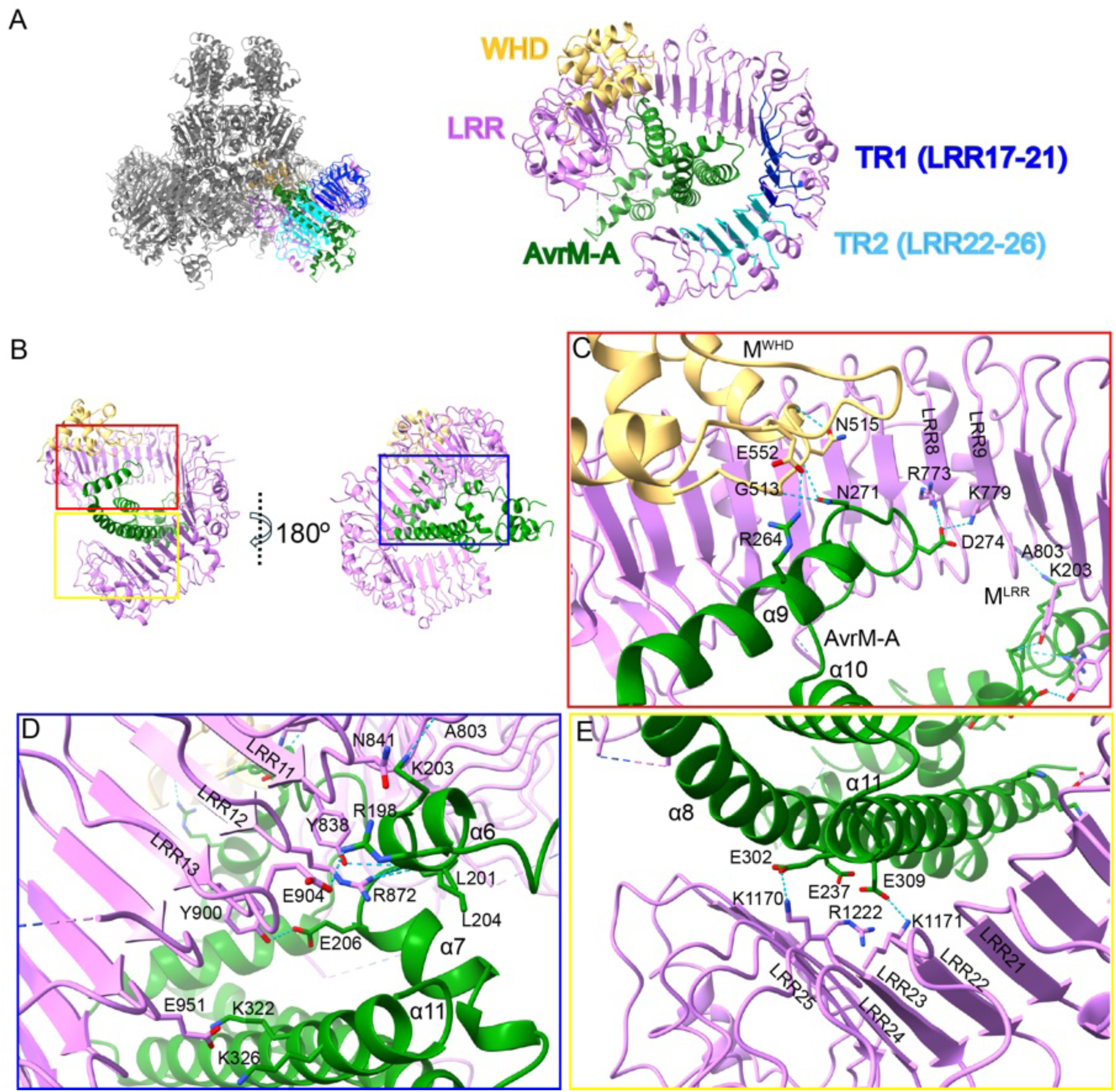
M directly interacts with AvrM-A protein via the WHD and LRR domains through polar interactions. **(A)** The M^WHD-LRR^:AvrM region of the M resistosome cryo-EM structure is highlighted (left panel) and shown in cartoon representation (right panel; TIR, NBD, and HD1 are not shown for clarity). The different domains/regions are colored: M^WHD^, yellow; M^LRR^, magenta; M^LRR^ TR1, blue; M^LRR^ TR2, cyan; and AvrM-A, green. **(B)** Structure of M binding to AvrM-A. The cryo-EM structure of M^WHD^ (yellow) and M^LRR^ (magenta) that interact with a monomer of AvrM-A (green) are shown in cartoon representation. **(C-E)** Close-up views of the interactions between M and AvrM-A in three different regions. The protein backbone is shown in cartoon representation; the coordinating residues are shown in stick representation and labeled. **(C)** Red box: two loops in the WHD and LRR8-9 motifs of M contact the α9-α10 region of AvrM-A. **(D)** Blue box: multiple LRR motifs of M interact with the α6-α7 loop and the α11 helix of AvrM-A. **(E)** Yellow box: interactions between TR2 of M^LRR^ and the α8 and α11 helices of AvrM-A, involving charged residues.

### The WHD and LRR domains of M contribute to effector binding

Superposition of the AvrM-A crystal structure (PDB: 4BJN) onto the AvrM-A monomer in the M resistosome showed that there are no large conformational changes in the effector upon recognition by M (Figure S5B). There are two domains of M (WHD and LRR) that directly interact with AvrM-A, mainly through polar interactions (Figure 3). The barley MLA13 similarly employs the WHD and LRR domains for the effector AVR_A13_ binding ^38^. In the WHD:AvrM-A interacting region (Figure 3B, C), two loops within M^WHD^ (residues 511-517 and 540-555; M^G513^, M^N515^ and M^E552^) contact the α9 helix and the α9-α10 loop of AvrM-A (AvrM-A^R264^ and AvrM-A^N271^) (Figure 3C). Interactions between the LRR region and AvrM-A involve multiple M^LRR^ motifs (Figure 3B-E). M^LRR8-9^ (M^R773^ and M^K799^) interact with AvrM-A^D274^ in the α9-α10 loop (Figure 3C). When compared with the superimposed AvrM-A crystal structure, this loop of AvrM-A in the cryo-EM structure is rotated further away from the main body of AvrM-A, allowing it to contact M^LRR^ (PDB: 4BJN; Figure S5B). More M^LRR^ motifs (LRR9, 11, 12, 13 and 15; M^A803^, M^Y838^, M^N841^, M^R872^, M^Y900^, M^E904^, and M^E951^) interact with the effector through the α6 helix (AvrM-A^R198^, AvrM-A^L201^, and AvrM-A^K203^), the α6-α7 loop (AvrM-A^L204^ and AvrM-A^E206^), and the α11 helix (AvrM-A^K322^ and AvrM-A^K326^), through polar interactions (Figure 3D). Furthermore, the TR2 region of M^LRR^ (LRR23 and 25; M^K1170^, M^K1171^, and M^R1222^) contacts the effector via helices α8 (AvrM-A^E237^) and α11 (AvrM-A^E302^ and AvrM-A^E309^) (Figure 3E). Consistent with our structural observations, previous studies demonstrated that the C-terminal truncation from the α8 helix (lacking residues 225-343) and the α11 helix (lacking residues 312-343) of AvrM-A abolished M-dependent cell death in *N. tabacum* ^32^, suggesting that this region is essential for M-mediated recognition.

### Dissociation of the AvrM-A dimers into monomers during recognition by M

The AvrM-A effector forms a stable homodimer in plants, solution and crystals ^32,35^. However, our cryo-EM structure revealed that each M protein interacts with a monomer of AvrM-A (Figure 2, 3). Superposition of the AvrM-A dimer crystal structure (PDB: 4BJN) onto the cryo-EM structure revealed that the second AvrM-A molecule would sterically clash with the TR2 region of M^LRR^ (Figure S5C). This observation suggests that the AvrM-A dimerization interface is involved in M^LRR^ interaction. The total buried surface area of the M:AvrM-A interface is greater (∼2,270 Å^2^) than that of the AvrM-A dimerization interface (∼1,950 Å^2^) ^35^. We reason that during recognition of AvrM-A by M, the AvrM-A dimeric protein dissociates into monomers. This mechanism resembles a previous observation that monomers of the dimeric stem rust effector AvrSr35 bind to each LRR domain of pentameric wheat CNL Sr35 ^9^. Based on the AvrM-A crystal structure, there are three charged residues (E237, E309 and R313) that form salt bridges in the AvrM-A dimerization interface (Figure 4A). Among these, the two glutamate residues also contribute to interaction with M in our M resistosome cryo-EM structure (Figure 3E). The AvrM-A protein carrying E237A, E309A and R313A triple mutations still retained the dimeric form (∼58 kDa) *in vitro*, resembling the wild-type AvrM-A protein behavior (Figure S6). This finding suggests that these amino acid substitutions are insufficient to disrupt effector dimerization, likely due to involvement of multiple other residues in the large dimerization interface. However, when the AvrM-A^E237A/E309A/R313A^ protein was incubated with M, we no longer detected formation of the M resistosome (Figure 4B). Consistent with this, the AvrM-A protein with the double E237A and E309A substitutions lost interaction with the M receptor in Y2H assays (Figure S7A), despite the protein being expressed at similar levels in yeast as wild-type AvrM-A (Figure S7B). Transient expression of AvrM-A carrying the double (AvrM-A^E237A/E309A^) or triple (AvrM-A^E237A/E309A/R313A^) mutations abrogated HR in *M*-transgenic *N. tabacum* leaves (Figure 4C), while single mutants or double mutants (AvrM-A^E309A/R313A^ and AvrM-A^E237A/R313A^) elicited M-dependent HR (Figure 4C). Collectively, our data indicate that both E237 and E309 of AvrM-A contribute to M recognition and immunity activation.

**Figure 4.**
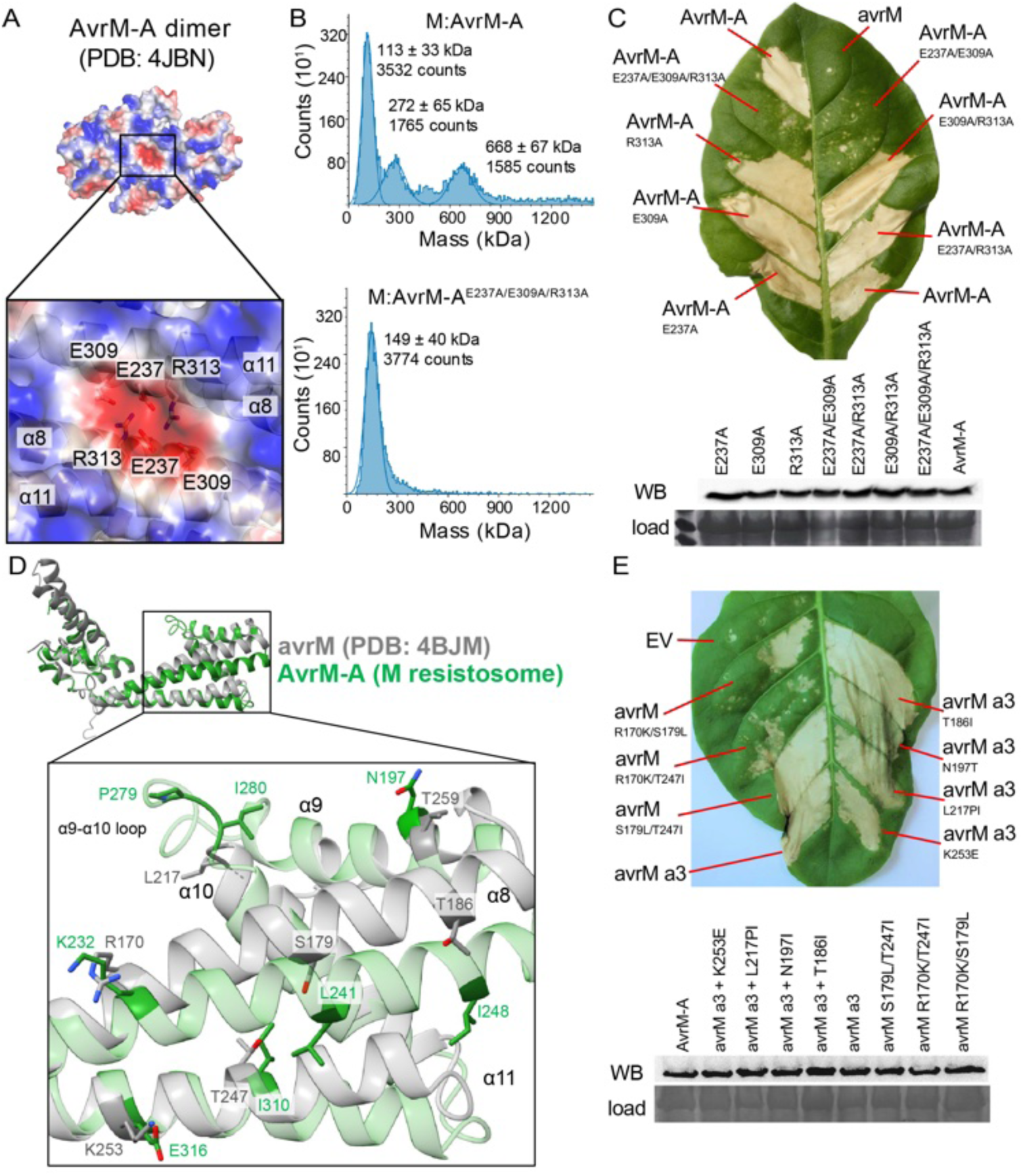
Involvement of effector residues in the α8 and α11 helices in receptor binding and recognition escape. **(A)** Crystal structure of the AvrM-A dimer (PDB: 4BJN) shown in surface representation and close-up view, showing charged residues (E237, E309 and R313) that form salt bridges in its dimerization interface. The secondary structures at the interface are shown in cartoon representation. The coordinating residues are shown in stick representation and labeled. **(B)** Mass photometry analysis of samples incubated with the monomeric full-length M protein, the wild-type (top) or mutant AvrM-A protein (bottom; E237A/E309A/R313A), ATP and 1AD. Upon M:AvrM-A incubation (top), peaks corresponding to the M monomer, dimer and tetramer were detected, while incubation with the AvrM-A mutant protein (bottom) did not form the M dimer and tetramer peaks. **(C)** Effects of transient expression of AvrM-A effector variants on M-dependent cell death and protein accumulation in leaves of *M*-transgenic *N. tabacum*. Effectors tested include the wild-type avirulence AvrM-A, the virulence avrM, and mutant AvrM-A variants carrying single, double and triple mutations, as indicated in the figure. **(D)** Structural superposition of virulence avrM (PDB: 4BJM, chain A) onto the avirulence AvrM-A protein in the cryo-EM structure of the M resistosome (top) and its close-up view of region from α8 to α11 helices (bottom). **(E)** Analysis of HR triggered by avrM variants transiently expressed in *M*-transgenic *N. tabacum*. Tested avrM variant samples include avrM ‘a3’ (R170K/S179L/T247I), double mutants (R170K/S179L, R170K/T247I, or S179L/T247I), and avrM a3+ variants (a3+K186I, a3+N197T, a3+L218PI or a3+K253E). **(C, E)** Protein expression is validated by western blotting (WB). Equivalent sample loading (load) is shown by SDS-PAGE.

### Mechanism by which the avrM effector escapes M recognition

The effector variant avrM, encoded by the virulence allele of this locus in flax rust, does not interact with M ^32^ and does not cause HR ^31^. AvrM-A and avrM (PDB: 4BJM) are structurally alike, with an RMSD (root mean square deviation) value of 1.48 Å for 185 Cα atoms ^35^. There are 13 polymorphic residues between AvrM-A (residues 108-343) and avrM (residues 46-280) and most of the AvrM-A residues that interact with M are identical in avrM (Figure S8A). We superimposed the avrM crystal structure (PDB: 4BJM) onto the AvrM-A within the M resistosome and observed that the α8 and α11 helices of avrM would position further away from M^LRR^, compared to those of AvrM-A involved in M-mediated recognition (Figure S8B). Three polymorphic residues (avrM^R170^, avrM^S179^ and avrM^T247^) lie within the α8 and α11 helices (Figure 4D; Figure S8A). Furthermore, the α9-α10 loop region of avrM is shorter than in AvrM-A and lacks a proline residue that is present in AvrM-A (Figure 4D; Figure S8A). To test if these differences in avrM account for the evasion of M-mediated recognition, we studied two avrM mutants designed to mimic AvrM-A. The first variant (designated the “a3” mutant; avrM^R170K/S179L/T247I^) carried three amino-acid substitutions, whereas the second contained an additional L217I substitution and an extra proline residue prior to I217 (designated “a3+L217PI”). SEC-MALS (size-exclusion chromatography coupled with multi-angle light scattering) analysis of the purified avrM variants, in the absence of the M receptor, showed that while avrM was present as a monomeric form (∼28 kDa), the avrM variants a3 and a3+L217PI avrM variants formed dimers resembling AvrM-A (∼55 kDa; Figure S6). We then evaluated the ability of the avrM mutants to interact with M. As expected, Y2H assays gave a positive interaction between M and AvrM-A, but no interaction was detected between M and the avrM mutant variants, although the protein expression was confirmed by immuno-blot analysis (Figure S7). The lack of detectable Y2H positive interaction between M and avrM mutant variants could be due to weak or transient interaction, combined with the stringency of the assayed conditions. Finally, transient expression of avrM a3 in leaves of *N. tabacum* led to elicitation of M-dependent HR in some, but not all the leaf infiltration assays (Figure 4E; Figure S9). However, the avrM a3+L217PI mutant, as well as avrM a3 carrying additional substitutions, including T186I, N197T or K253E located on the α8, α9 and α11 helix, respectively, consistently triggered HR (Figure 4E; Figure S9). Overall, our findings suggest that the mutations within the α8, α9 and α11 helices and the α9-α10 loop of avrM contribute to its escaping recognition by M, by possibly altering the orientation of the helices and the loop, disrupting interaction with M.

### The NB-ARC domain of the M resistosome is in the ADP-bound state

The NB-ARC domains of M assemble into a tetramer in a similar manner as those of RPP1 and ROQ1 (Figure 5A). While it is generally accepted that ATP binding induces oligomerization of NLRs, an ADP molecule, unambiguously defined by the cryo-EM density, is bound to the NBD in the M resistosome (Figure S10A). There was also additional EM density allowing us to model an Mg^2+^ ion (Figure S10A). Generally, the reported plant NLR resistosome structures are bound to ATP (or dATP) ^7–11,16^, except for the ADP-bound RPP1 tetramer ^15^. Based on structure comparisons, we confirmed that the nucleotide binding pocket of M^NBD^ involves the highly conserved motifs, including the P-loop (G-M-G-G-I-G-K-T-T) involved in nucleotide binding, the Walker-B motif (I-L-V-V-L-D-D) involved in ATP hydrolysis, and the sensor-1 motif (T-S-R) involved in ATP binding (Figure 5B). We observed that the P-loop residues directly interact with the β-phosphate (M^G283^, M^G285^, M^K286^, and M^T287^) and α-phosphate (M^T288^) moieties of ADP (Figure 5B). In addition, the N-terminal loop of the NBD (M^D250^ and M^V253^) interacts with a ribose and an adenosine group, and the GLPL motif in HD1 supports the positioning of ADP ^39^, as observed for other structures (Figure 5B; S10B). Comparison of the nucleotide binding region of TNL resistosomes showed some differences. M has a positively-charged arginine residue (R392) within the sensor-1 motif of NBD, which is required for contacting the γ-phosphate moiety of ATP for ROQ1 (Figure 5B, S10B), as well as the animal Apaf-1 (apoptotic protease activating factor-1) apoptosome and CNL resistosomes, suggesting that M can bind to ATP. By contrast, many other TNLs, including RPP1, instead have a negatively-charged glutamate in this motif ^15^ (Figure S10B). Furthermore, RPP1 has an extended β2-α2 loop (residues 320-331) that uniquely contributes to ADP-binding, by interacting with the P-loop of the adjacent RPP1 molecule ^15^. The analogous loop of M is shorter (Q-E-Q-K-D-G) (Figure 5A), and hence, it is unlikely to be involved in the stabilization of the ADP-bound form. Additionally, we found that M possesses an arginine residue (R488) in WHD, which forms hydrogen bonds with α- and β-phosphate groups of ADP (Figure 5B, Figure S10). Such an arginine residue is absent in all other reported plant NLR resistosomes. Therefore, both the conserved motifs as well as the M-specific residue contribute to stabilizing the ADP-bound state.

**Figure 5.**
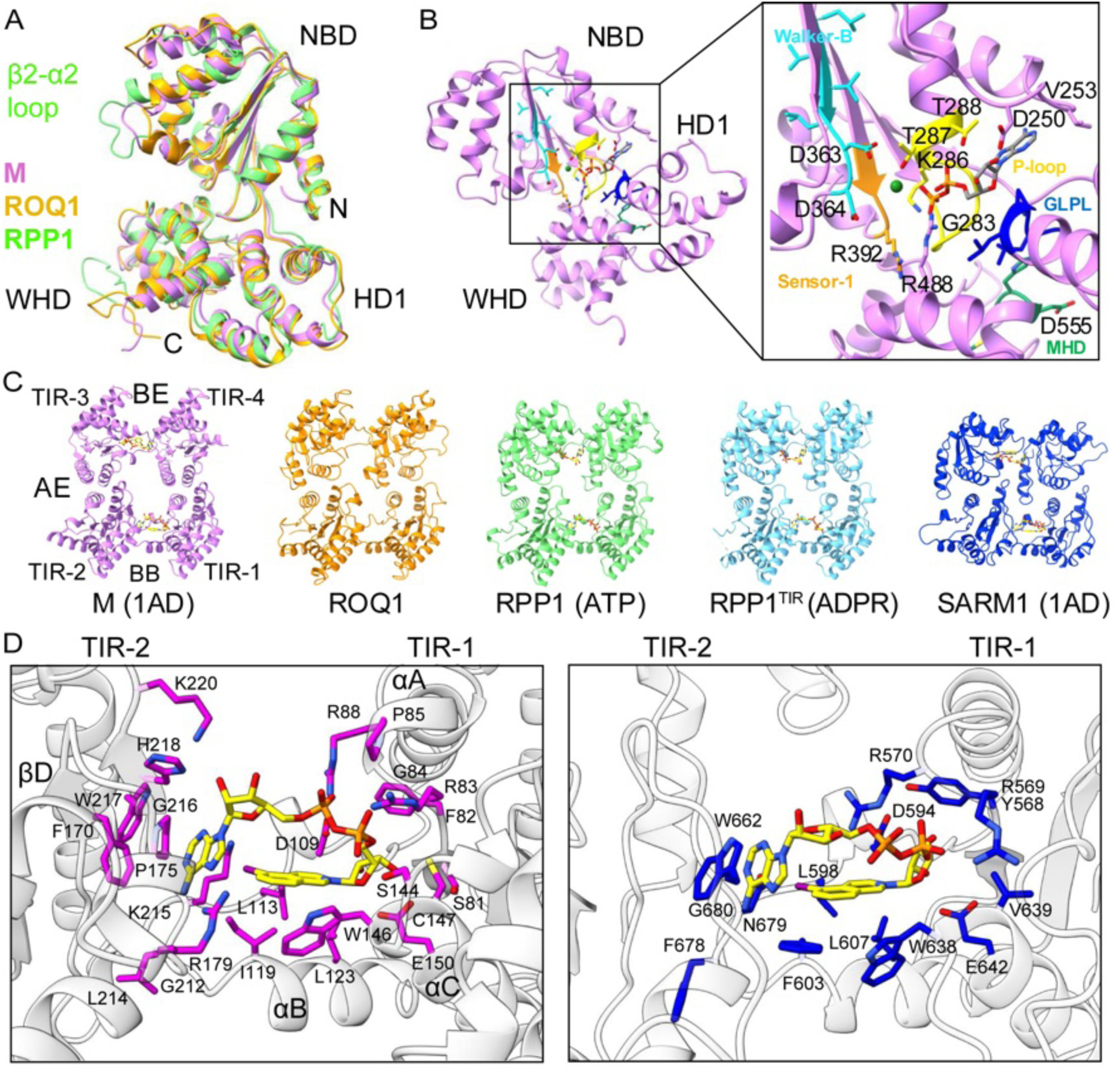
Ligand binding by the NB-ARC and TIR domains of M in its activated state. **(A)** Structural superposition of the NB-ARC domain of M, ROQ1 and RPP1 TNL resistosomes. M, ROQ1 and RPP1 are shown in purple, orange and green, respectively. Subdomains of the NB-ARC domain are indicated: NBD (nucleotide binding domain), HD1 (helical domain 1) and WHD (winged-helix domain). The β2-α2 loop is extended in RPP1 compared to that of M and ROQ1. **(B)** ADP is bound to the NB-ARC domain of the M resistosome. Conserved motifs for ADP/ATP binding are shown in the close-up view (right), including the P-loop (yellow), Walker-B (cyan), sensor-1 (orange), GLPL (blue), and MHD (green). The Mg^2+^ ion is shown (green) motifs. **(C)** Comparison of TIR domain assemblies of M (1AD-bound), ROQ1 (no ligand; PDB: 7JLX), RPP1 (ADP-bound; PDB: 7CRC), RPP1^TIR^ (ADPR-bound; PDB: 7XOZ) and SARM1 (1AD-bound; PDB: 7NAK), representing the activated state. Two symmetric TIR homodimers formed through the AE interface further associate with each other, forming the BE interface. The TIR dimer/two-fold symmetry is different to the four-fold symmetry of the rest of the complex. **(D)** 1AD binds to the catalytic pocket of the M resistosome (left panel) and human SARM1 (right panel).

### The TIR domain assembly and the NAD^+^ substrate recognition

TIR domains of the M resistosome form two symmetric (head-to-head) homodimers associated through the AE interface (αA and αE helices and their neighbouring loops; Figure 5C). These two TIR homodimers pack against each other, resulting in an asymmetric (head-to-tail) interaction. This interaction is referred to as the BE interface, where the BB-loop (between the βB strand and αB helix) of one TIR molecule is positioned under the DE loop (between the αD_3_ and αE_1_ helices) of an adjacent TIR molecule; here is where the substrate NAD^+^ binding site forms (Figure 5C).

We compared the AE interface of flax M with that of previously reported structures, including *Muscadinia rotundifolia* RPV1, *Vitis rotundifolia* RUN1, Arabidopsis RPP1 and *N. benthamiana* ROQ1 (Figure S11A). Superposition onto the TIR-1 molecule of M shows that the relative position of M TIR-4 reaches closer towards its TIR-1 molecule, making the AE interface of M more compact compared to other structures. From previous structural and functional studies on plant TIR domains ^14,36,40,41^, the core of the AE interface involves (i) a conserved histidine or phenylalanine residue from αA, which forms a stacking interaction between two TIR monomers, and (ii) an adjacent aspartic acid or serine residue from αA, forming hydrogen bonds with residues from αE. These residues are required for triggering cell death in plants. When comparing the AE interface between RPV1^TIR^ ^40^ and M^TIR^, instead of a histidine or phenylalanine, M has an isoleucine (I94), while the aspartic acid residue (D93) is conserved in RPV1 (D41; Figure S11, S12). Substitution of isoleucine and/or the adjacent aspartic acid with alanine (I94A and/or D93A) abrogated AvrM-A-triggered cell death in *N. benthamiana* (Figure S13A). These mutant proteins were similarly expressed as wild-type M (Figure S13B), suggesting that both residues are key to forming the AE interface of M. M further has several polar interactions that differ from previously reported structures; D230 (αE) contacts R97 and R101 (αA) (Figure S11B), suggesting that these charged residues also contribute to stabilizing the AE interface.

In the catalytic pockets formed within the BE interface, the ROQ1 or RPP1 resistosome structures either have no substrate or an ATP molecule modelled into the EM density, respectively ^15,16^ (Figure 5C). On the other hand, a crystal structure of the TIR domains of RPP1 was shown to bind to ADPR (ADP ribose) when supplemented with NAD^+^ ^22^, suggesting that NAD^+^ was cleaved into nicotinamide and ADPR (Figure 5C; Figure S13C). Our sample included the NAD^+^ mimetic 1AD, which is clearly observed bound to the TIR catalytic sites; the BB-loop of the TIR-1 molecule reaches towards the DE surface of the TIR-2 molecule (Figure 5C, D; Figure S14A). 1AD is a base-exchange product, resulting from the replacement of the nicotinamide group of NAD^+^ with compound “1” (5-iodo-isoquinoline) ^34^. M^TIR^ binds 1AD in an analogous manner as human SARM1^TIR^ ^34^ (Figure 5D). In the TIR-1 molecule, two highly conserved arginine residues (M^R83^ and M^R88^), N-terminal to the αA helix (not part of the AE interface), have side chains pointing towards the substrate-binding site on top of the BB-loop, and form hydrogen bonds with the diphosphate groups of 1AD (Figure 5D). The arginine residue, equivalent to M^R88^, has been proposed to play a key role in opening the BB-loop, to create the NAD^+^ binding site upon activation ^42^. Indeed, substitution of this arginine to alanine in RUN1 (R39A) resulted in loss of *in vitro* NADase activity ^42^, and similar mutations in L6 (R73A) and RPV1 (R36A) abolished HR ^40,43^. Three residues, M^I119^ (BB-loop), M^L123^ (αB), and M^W146^ (αC), stabilize the aromatic rings of the 5-iodo-isoquinoline group, analogous to SARM1^TIR^ residues (F603, L607 and W638). The conserved M^E150^ (αC) forms a hydrogen bond with the C-3 hydroxyl group of the ribose. Next, in the TIR-2 molecule, the conserved phenylalanine M^F170^ (βD) stacks against the adenine group, resembling SARM1^W662^. As described for SARM1^N679^, M^K215^ forms a halogen bond with the 5-iodo group of 1AD. In contrast to SARM1^TIR^:1AD, we observed involvement of additional M amino-acid residues engaging in ligand binding (Figure 5D). The diphosphate groups additionally form hydrogen bonds with the highly conserved residues M^G84^ (prior to αA) and M^S144^ (αC) of TIR-1. M^H218^ in TIR-2 also forms a hydrogen bond with the adenine-linked ribose group. Multiple residues in the DE surface of TIR-2 (M^R179^, M^G212^, M^L214^, and M^G216^) interact with the adenine group by hydrogen bonding; they are stabilized by surrounding residues (M^P175^, M^K220^ and M^K215^). Multiple residues analogous to the ones in M are involved in the interaction with the ADPR in the RPP1^TIR^:ADPR crystal structure (Figure S13C).

To corroborate the analysis above, we introduced mutations targeting the two interfaces and tested their effects on the ability to trigger HR in *N. benthamiana*, compared to wild-type M (Figure S13A, B). Single amino-acid substitutions of highly conserved residues (the catalytic glutamate (M^E150A^) in the αC helix, and glycine (M^G216R^) and adjacent tryptophan (M^W217A^) in the αD_3_-βE loop, which presumably disrupt interactions with the ribose and adenine group of NAD^+^, respectively), abolished effector-dependent cell death (Figure S13A). Consistently, the impaired cell death phenotype was observed in previous studies when mutations similar to M^E150A^ (L6^E135A^, SNC1^E84A^, RPS4^E88A^, RUN1^E100A^, RBA1^E86A^, BdTIR^E127A^, RPP1^E158A^, ROQ1^E86A^), M^G216R^ (L6^G201R^, RPV1^G161R^, ROQ1^G153A^), and M^W217A^ (L6^W202A^, SNC1^Y150A^) ^14–17,41,43^ were tested. On the other hand, substitution in the conserved tryptophan (M^W146A^) or serine residue (M^S144A^) within the αC helix was insufficient to attenuate AvrM-induced HR (Figure S13A), likely due to multiple other neighbouring residues stabilizing the NAD^+^ substrate. Similar substitutions of other plant NLRs resulted in variable phenotypes. For instance, effector-dependent HR was completely (for L6^S129A^) or partially (for N^W82A/S^) abrogated ^43,44^. TIR domain autoactivity was impaired in the L6^W131A^ and RPV1^W93A^ mutants ^40,43^, whereas, by contrast, autoactivity of RPS4^W84A^ was enhanced ^45^. Mutations in residues that are part of the BE interface but do not contribute to direct substrate interaction (M^K176E/A^ and M^K215E^), triggered HR similar to wild-type M (Figure S13A). All the mutant proteins tested above were detected by western blot, but an additional mutant (M^K145A^ in the αC helix) did not accumulate (Figure S13B). Overall, findings from our *in planta* assays and earlier studies support our structural observation that multiple TIR residues are involved in formation of the BE interface and binding to the NAD^+^ substrate.

### NADase activity by flax M and L6 TNLs leads to the production of 2’cADPR and ADPr-ATP

We next characterized the NADase activity of full-length flax M and the closely related TNL L6. L6 was expressed and purified using similar approaches as described for M. Using fluorescence-based NADase assays, we detected that both M and L6, at a concentration as low as 300 nM, readily cleaved the fluorescent NAD^+^ analogue εNAD (1,N^6^-ethenoNAD), in the presence of 1 μM effector and 1 mM ATP (Figure S14B). No fluorescence was detected in the absence of the effector and/or ATP (Figure S14B), suggesting that NADase activity relies on the resistosome formation. We next evaluated effects of 1AD on NADase activity of M and L6. Incubation of NLR, effector and ATP with 1AD prior to adding εNAD resulted in no fluorescence increase (Figure S14B), suggesting that 1AD is indeed an inhibitor of NAD^+^-cleavage activity by plant TNLs.

Previous studies revealed that NAD^+^ cleavage by the TIR domain of L6 leads to the production of 2’cADPR ^17,46^. Here, we sought to identify small molecule products generated upon NADase activity by both M and L6 full-length proteins, upon effector-induced activation *in vitro*. We confirmed by 1D ^1^H NMR (nuclear magnetic resonance) that only when cognate effectors and ATP were supplied in the reaction mixture, M and L6 cleaved NAD^+^ and produced 2’cADPR, albeit at low levels (indicated by a small peak; Figure 6A; Figure S15). Through MALDI-TOF (matrix-assisted laser desorption/ionization – time-of-flight) mass spectrometry analysis, we also detected a chemical species corresponding to the mass of ADPr-ATP + 1H (*m/z* = 1049.06) by activated M and L6 (Figure 6B; Figure S16). We did not observe any signals for 3’cADPR and di-ADPR. No catalytic activity and products were detected when the reaction mixture lacked ATP and/or the effector (Figure 6; Figure S15, S16). In summary, flax rust effector-activated M and L6 function as NADases and their enzymatic activity leads to generation of a common set of nucleotide-derived products, 2’cADPR and ADPr-ATP.

**Figure 6.**
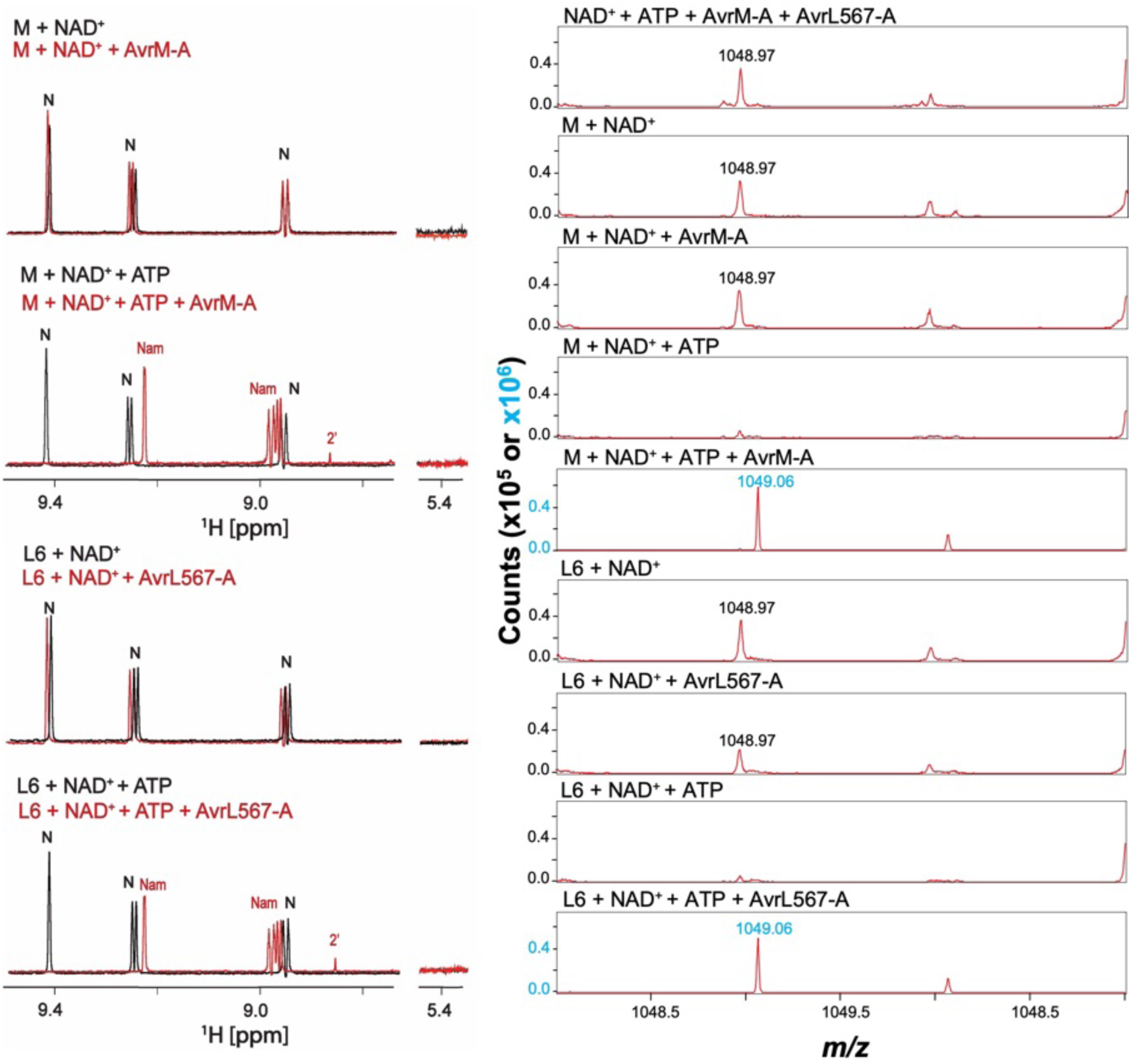
Catalytic products derived from NADase activity by activated flax M and L6 receptors. **(A)** ^1^H NMR spectrum from 9.5 – 8.75 ppm and at 5.4 ppm, showing M and L6 incubated with NAD^+^ or NAD^+^ with ATP together, with and without their cognate effectors. Peaks are labeled to show which compound they represent, N (NAD^+^), Nam (nicotinamide) and 2’ (2’cADPR). Labels are color-coded black (sample without effector) and red (with effector), to show which sample they relate to. In samples without ATP, only NAD^+^ is identified, whether the effector was added or not. The 5.4 ppm region is where a peak corresponding to 3’cADPR would be expected, but it was not present in any of the tested samples. An extended spectrum with the inclusion of standards is shown in Figure S15. **(B)** MALDI-TOF MS analysis showing *m/z* spanning from 1048 – 1051 *m/z.* The black label indicates a background species present at 0 h and therefore is not related to catalytic products; the while blue label (1049.06) indicates *m/z* corresponding to ADPr-ATP. The y-axis (counts) was decreased 10-fold for samples without NAD^+^ cleavage, to show no background of the 1049.06 *m/z* species. MS/MS analysis is shown in Figure S16.

### The monomeric M receptor has a closed conformation

In the absence of pathogens, NLRs are considered to stay in a closed conformation mediated through intramolecular interaction, thereby inhibiting their oligomerization and activation of immune signaling. Here, we collected cryo-EM data for the monomeric M protein and obtained an EM density map at an overall resolution of 6.9 Å (Figure 7A; Figure S17). Each individual domain, except for the TIR domain, of the M protomer from the resistosome could be docked into the cryo-EM map (Figure 7A). The absence of cryo-EM density for M^TIR^ suggests that this domain in the monomeric protein is flexible, which is unlike ZAR1^CC^, which interacts with its LRR, WHD and HD1 in the monomeric form ^6^. The AlphaFold 3 ^47^ predicted monomeric M protein structure showed a similar arrangement for the NB-ARC and LRR domains, whereas the TIR domain was positioned far above the lateral side of N-terminal LRR, facing the WHD side (Figure S18A). We found that NBD, HD1 and WHD of the M monomer model (without the TIR domain) are positioned similarly to those of the autoinhibited state of CNLs, including Arabidopsis ZAR1 and *Solanum lycopersicum* NRC2 (Figure S18B). Furthermore, the positioning of the LRR domain relative to the NB-ARC domain of these plant NLRs is the same (Figure S18B), whereas in animal NLRC4^LRR^, it is oriented differently ^6,48^. M has more LRR motifs than ZAR1 and NRC2 proteins, and the M monomeric protein is in a closed conformation where M^NBD^ is positioned on top of the lateral surface in the middle of M^LRR^, thereby capping the LRR domain (Figure 7; Figure S18).

**Figure 7.**
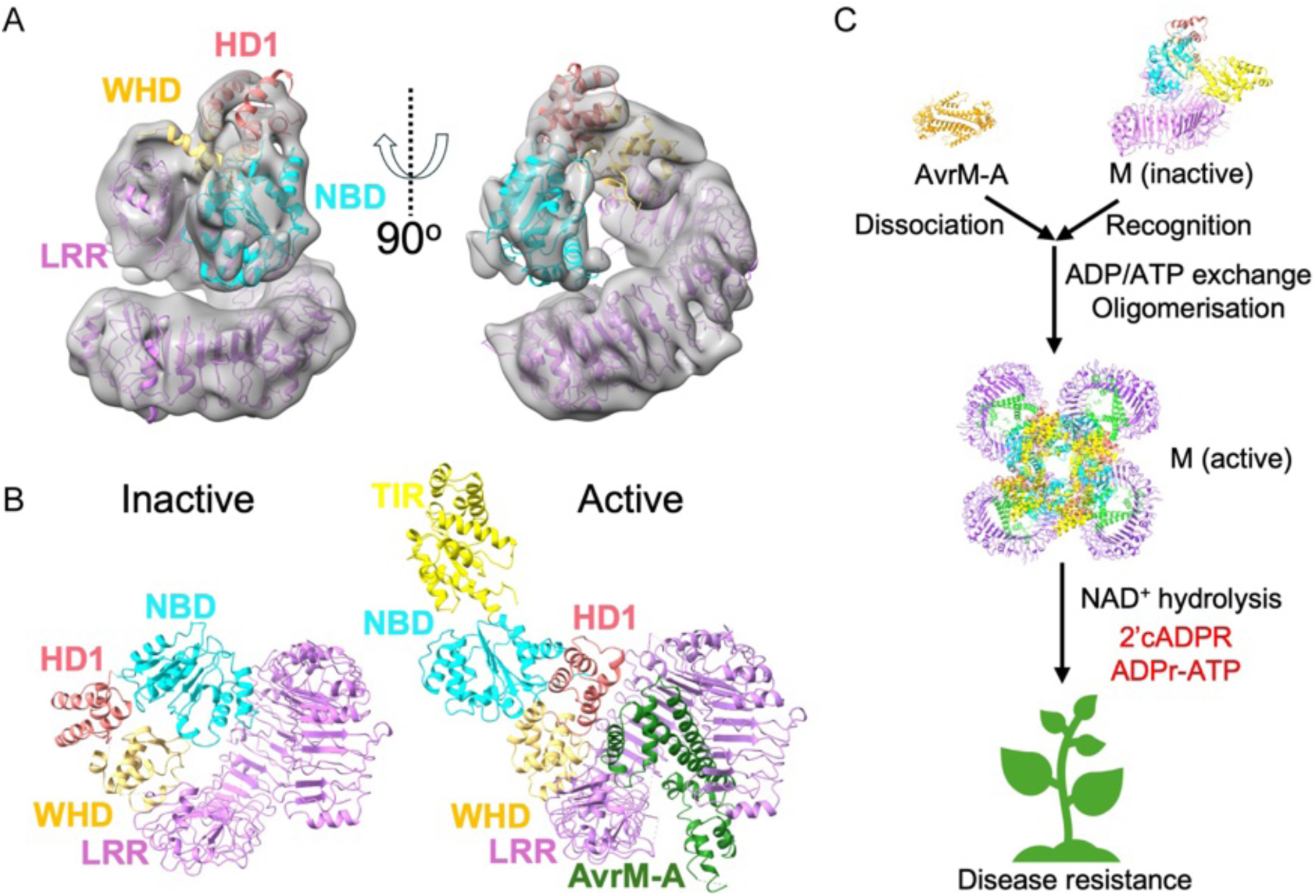
The M monomer structure represents the inactive state. **(A)** Cryo-EM reconstruction of the M monomer. Each NBD, HD1, WHD and LRR domain from the protomer of M from the resistosome model was fitted into the EM density. The EM density for the TIR domain is absent. **(B)** Structural comparison between the inactive M monomer and the active M (a protomer from the M resistosome) shows different positioning of their NBD and HD1. **(C)** Working model for M activation induced by AvrM-A. The M monomer is in a closed conformation, preventing autoactivity. During AvrM-A recognition, the effector dimer dissociates into monomers and induces exchange of ADP for ATP in the NBD and a conformational change in the NB-ARC region, leading to formation of the M resistosome. The assembled TIR domains form two NAD^+^-binding pockets, resulting in NAD^+^-hydrolysis and formation of the small molecules 2’cADPR and ADPr-ATP, which act as signaling molecules that activate downstream immune pathways.

Structural superposition of the LRR domain between the AvrM-A-bound M protomer from the resistosome and the M monomer shows a steric clash between AvrM-A (α8-α9 and α10-α11 loop regions) and M^NBD^ (Figure S18C). The positioning of M^NBD^ and M^HD1^ domains is substantially different between the monomeric and resistosome models, because M^NBD^ would flip away from M^LRR^ to accommodate a monomer of AvrM-A, and M^HD1^ would flip towards M^LRR^ to accommodate M^NBD^ (Figure 7B). The relative position of M^WHD^ of the M monomer appears to be similar to that of the resistosome (Figure 7B). In conclusion, our observations suggest that the M monomeric protein represents the inactive state with this autoinhibited conformation being conserved among different classes of plant NLRs; the structures are consistent with the conformational changes observed in the CNL ZAR1 when transitioning from inactive to active states ^6,7^.

## Discussion

In this work, we report the cryo-EM structures of a canonical TIR-domain containing NLR, with a prototypical tripartite domain architecture, in its resting state and effector-activated state. The mechanism of effector recognition and activation of the flax M receptor involves the following steps: (i) Upon infection by *M. lini*, monomeric M, in its autoinhibited state, detects the AvrM-A dimeric effector, leading to AvrM-A dimer dissociation and direct interaction of its monomers with the WHD and LRR domains of M; (ii) effector recognition induces the M protein NBD and HD1 domain rearrangement and exchange of ADP for ATP on the NB-ARC domain, followed by NLR oligomerization into dimers (intermediate form) and tetramers (the resistosome); and (iii) the assembled TIR domains of the M resistosome create two NAD^+^-binding pockets, resulting in NAD^+^ cleavage and generation of second messengers, 2’cADPR and ADPr-ATP, that activate downstream immune signaling and ultimately disease resistance (Figure 7C).

Our cryo-EM analysis of the M monomer suggests that it forms an autoinhibited state, in agreement with the previously observed NB-ARC domain arrangement of CNLs ^4–6^. Therefore, both TNLs and CNLs share a conserved closed conformation in their resting state. Interestingly, while inactive ZAR1 CNL is a monomer, inactive helper NRC2 proteins from *N. benthamiana* and *S. lycopersicum* form homodimers ^4,5^ that can further assemble into tetramers or filaments, depending on sample concentrations *in vitro* ^4^. Such higher-order assemblies are reported to be required to stabilize the autoinhibited state *in planta* ^4^. It is notable that in our SEC experiments, the purified M protein was eluted over a range of elution volumes (Figure 1A, B), consistent with previous studies ^49^. These observations point out that in the absence of AvrM-A, the M receptor *in vitro* may form a mixture of inactive monomers, inactive higher-order oligomers and/or possibly non-functional aggregated forms (Figure 1A, B). Future studies should address whether M or other NLRs form a multimeric autoinhibited complex, reminiscent to or distinct from NRC2 oligomers, and if so, its physiological relevance.

The M resistosome cryo-EM structure revealed that, in the LRR domain, the second tandem repeat (TR2) region is crucial for interaction with AvrM-A. Because this tandem repeat region would sterically clash with the effector dimerization interface, it appears to serve as a site for competitive binding and enforcement of dissociation of the dimeric effector into monomers, as part of receptor activation. Diversification of the LRR tandem repeat regions by duplication can be one of the factors driving novel effector recognition specificities. Consistent with our findings, earlier work identified that three *M* mutant alleles lacking one of the tandem repeats lost *M* resistance specificity to flax rust fungi ^30^. These mutant alleles likely arose from unequal intragenic recombination events ^30^. Furthermore, the related flax *M1* gene encodes a TNL with three tandem repeat copies of sets of LRRs ^50^ and recognizes a distinct effector allele *AvrM14* encoding a nudix hydrolase ^51,52^. Another example is the flax *L* gene with 13 allelic variants, exhibiting distinct recognition specificities. In the *L1*, *L2* and *L8* alleles, major differences are found in the sequences encoding the C-terminal LRR domain, with loss of the second and first tandem repeat copy in *L1* and *L8*, respectively, and presence of four copies in *L2* ^53^. *L2* is known to recognize *AvrL2*, which encodes a highly charged, small proline-rich protein; the LRR domain seems to be fundamental for *L2* specificity ^51^.

In the resistosome structure, we observed that M^NB-ARC^ was bound to ADP (Figure 5A, B; Figure S10), despite being supplemented with 1 mM ATP in the reaction mixture. This could implicate its post-activation state, where ATP had undergone hydrolysis to ADP. The wild-type M protein, presumably in its resting state without the effector, binds to ADP *in vitro*, whereas the autoactive M^D555V^ mutant displays preferential binding to ATP and triggers effector-independent HR *in planta* ^49^. These findings suggest that M probably exchanges ADP for ATP upon activation. We also used prolonged incubation time for formation of the M resistosome prior to plunge-freezing, during which ATP hydrolysis could have occurred. It remains to be elucidated whether ADP binding is sufficient to induce the formation of resistosomes, particularly for those NLRs, such as RPP1, that lack the conserved sensor-1 and Walker-B motifs.

M^TIR^ binding to the NAD^+^-mimetic 1AD revealed that multiple residues in the DE surface, the BB-loop and the αC helix are essential for the NAD^+^ substrate binding. In our *in vitro* NADase assays, we further demonstrated that both M and L6 cleave NAD^+^, upon incubation with their cognate effectors and ATP, leading to ADPr-ATP and 2’cADPR production. We also detected other compounds, corresponding to *m/z* = 462.1 and *m/z* = 638.1 (Figure S16C), consistent with additional molecules produced by several other activated TIR domains ^54^; they may be degradation products of ADPr-ATP, di-ADPR or pRib-ADP. However, their identity and biological relevance remain to be addressed in future studies. It is yet to be established how the products detected *in vitro* relate to products *in planta*, as well as the outcomes of the immune response. In Arabidopsis, ADPr-ATP can be captured in the conserved ligand binding pocket of the EDS1:SAG101 heterodimer ^22^, which then recruits the helper NLR NRG1 to activate immune signaling ^18,19^. 2’cADPR has been considered a marker of plant immune signaling but is insufficient to directly initiate EDS1-dependent signaling ^17,46,55^. In our *in vitro* assays, only a small amount of 2’cADPR was detected. The amounts of signaling compounds produced in plants could also be variable. Differences were indeed observed for *Arabidopsis* TIR domains, where among ∼70 TIRs that cause cell death in an EDS1-dependent manner, only ∼20% produced 2’cADPR at detectable levels in plants ^54^. Notably, 2’cADPR can also be produced by bacterial TIR domains, including the HopBY effector from *Pseudomonas syringae* that causes plant susceptibility ^56^. While the exact role of 2’cADPR in plants remains to be confirmed, recent studies show that 2’cADPR can be converted into pRib-AMP ^20^, which then activates the EDS1:PAD4 heterodimer to recruit the ADR1 helper NLR ^20,21,23^.

Despite the similarities between M and L6 in their TIR domain sequences and the signaling compounds they produce *in vitro*, we observed differences in cell-death phenotypes when some of the highly conserved residues were mutated (Figure S13B) ^43^. Furthermore, while transient expression of the TIR domain of L6 induces cell death *in planta* ^14,41,43^, that of M (and the related M1) did not, even when the SAM (sterile α-motif) domain of SARM1 was fused to force oligomerization of the proteins (Figure S19). These observations suggest that TIR activity and signaling output by these related TNLs are differentially regulated in plant cells. Previous studies found that genomic fragments of *M* and of several other TNL genes are required for conferring disease resistance, while their cDNAs were insufficient to trigger the immune response ^49,57,58^. These observations could be explained by alternative splicing, which has been implicated in regulating TNL expression and function ^57,58^. Expression of *M* leads to a complex transcript profile with 12 alternative isoforms derived from splicing at different introns (intron 1-2 and micro-open reading frame at 5’ UTR), most of which would translate into TIR and TIR-NB truncated products ^59^. By contrast, there are four *L6* alternative transcripts at intron 3, and transgenic flax plants with the full-length *L6* cDNA are capable of triggering a full resistance response ^60^. These findings suggest that, although the specific role of alternative splice variants of *M* remains to be established, they are likely essential to control M expression and function. At the same time, M and L6 localize to different subcellular compartments, tonoplast and Golgi, respectively ^61^. It is possible that yet unknown proteins or co-factors in specific cellular locations may influence the specific TNL enzymatic activity and the signalling molecules produced. A recent study reported that L6^TIR^ can elicit cell death when localized to Golgi or tonoplast, but not in the nucleus ^62^, and further investigation for M should provides insights. Understanding the TIR-derived compounds and their role *in planta* will also be important in developing efficient strategies for designing resistant crops. A recent study showed how TIR domain signaling output could be tuned by targeting the BB-loop ^54^. Building on these findings, future studies should explore and determine other key TIR residues required for product formation and variability. For example, the DE surface, in which the residues are variable among TIR domains, could also be an interesting region to test for altered amounts and types of products, as well as for effects on disease-resistance outcomes.

In conclusion, we present the first three-dimensional structures of a canonical TNL, with no C-JID domain, in both inactive monomeric and active resistosome states. Notably, the AvrM-A effector dimer dissociates upon binding to the M TNL. We visualize the binding of a non-hydrolyzable NAD^+^ analogue to the TIR-domain active site and show that the TNL produces the signaling molecules 2’cADPR and ADPr-ATP. Our findings explain the mechanism of canonical TNL signaling and provide a foundation for rational engineering of disease-resistant crops.

### Limitations of the study

While our cryo-EM anlaysis of M in its monomeric form revealed that it has a closed conformation representing the resting state, similar to that observed for the CC domain-containing plant NLRs, at this stage we were not able to obtain a high resolution structure and could not observe cryo-EM density for the bound nucleotide and the TIR domain. Future work on inactive TNLs should focus on optimizing parameters, including but not limited to, using ligands to potentially stabilize the inactive state and screening cryo-EM grids with additives and alternative buffer conditions to potentially reduce sample aggregation.

It is established that avrM encoded by the virulence allele exists as a monomer, while the avirulence AvrM-A forms a dimer, when tested in solution, by yeast two-hybrid assays and *in planta* transient expression followed by co-immunoprecipitation assays ^32^. However, only monomers of AvrM-A bind M, and the biological significance of the effector oligomeric status remains unclear. It is tempting to speculate that the effector dimerization interface may be functionally important and therefore highly conserved, which could prevent it from accumulating mutations to evade recognition by M. At the same time, we also cannot rule out the possibility that AvrM-A dimers could disassemble into monomers during translocation from the rust to the host. A recent study highlighted the importance of studying the effector delivery mechanism in the context of a natural infection process, by using transgenic rust lines expressing AvrM-A (with YFP fusion) under their native promoter and terminator, rather than transient overexpression in plants ^63^. Further comparative analysis of AvrM-A and avrM in their mechanism of translocation as well as identification of host target proteins are essential to provide insights into the effector biology and guide NLR engineering strategies.

## Acknowledgements

The work has been funded by the Australian Research Council (ARC; Discovery Project DP220102832 to BK and PND; Laureate Fellowship FL180100109 to BK; Future Fellowship FT200100572 to TV; Future Fellowship FT200100135 to SJW; Discovery Early Career Research Award DE130101292 to MB); the National Health and Medical Research Council (NHMRC Australia; Investigator grant 2025931 to BK, Investigator Grant 1196590 to TV); and the Swedish Research Council (2019-00815 to HX). We thank members of the Kobe laboratory group and colleagues (Yu Shang Low, Naphak Modhiran, Farrah Blades) for useful discussions. We thank the facilities and the scientific and technical assistance of SciLifeLab (Science for Life Laboratory, Sweden), Centre for Microscopy and Microanalysis (CMM) and Protein Expression Facility (PEF; University of Queensland, Australia). This work was undertaken with the assistance of UQ onBunya Open OnDemand and the EM Data Processing Portal. The EM Data Processing Portal is supported by the Australian Research Data Commons (ARDC), Monash University and the Queensland Cyber Infrastructure Foundation (QCIF). We thank the UQ Research Computing Centre and staff (Edan Scriven, David Green and Marlies Hankel) for technical support.

## AUTHOR CONTRIBUTIONS

Conceptualization

BK, JGE, PAA, SJW, PD

Data curation

NM, MS, YL, HX, LW, KK

Formal analysis

NM, WG, BYJL, MS, DHYN, CJ, JDN, JC, MB, SJW, PAA

Funding acquisition

HX, MB, TV, SJW, BK

Investigation

NM, WG, BYJL, MS, DHYN, JDN, YL, JC, MO, MB, MR, TDV, HX, LW, KK, SJW

Methodology

NM, WG, BYJL, MS, DHYN, CJ, JDN, JC, MO, HX, LW, KK, HB, SJW

Project administration

BK

Resources

HX, MM, MB, TV, BK

Supervision

MM, PAA, SJW, PD, BK

Validation

NM, WG, BYJL, MS, DHYN, CJ, JDN, JC

Visualization

NM, WG, BYJL, MS, DHYN, JDN, JC, MB, TDV

Writing – original draft

NM, BK

Writing – review & editing

All authors

## Resource availability Lead contact

Further information and requests for resources and reagents should be directed to and will be fulfilled by the lead contact, Bostjan Kobe.

## Materials availability

All the plasmids generated in this study are available from the lead contact.

## Data availability

The cryo-EM maps have been deposited in the EMDB with accession numbers EMD-71849 (M resistosome; consensus map), EMD-71848 (M resistosome; M^LRR:AvrM-A^ local map), EMD-71832 (M resistosome; composite map) and EMD-72188 (M monomer). The atomic models of the M resistosome and M monomer have been deposited in the PDB with accession numbers 9PT6 and 9Q38, respectively.

## Supplemental information

Figures S1–S19, Tables S1 and S2

## Materials and Methods

### Plasmid construction, protein expression and purification of full-length M

A DNA fragment encoding a Twin-Strep tag followed by a TEV (tobacco etch virus) protease cleavage site was synthesized and cloned into the previously reported construct, pPICZ carrying 6xHis-M (residues 21-1305) ^33^, at the *SacII* and *NotI* sites. Additionally, to reduce premature transcripts during expression in yeast cells, an AT-rich sequence found in the TIR domain-encoding region was modified from TTTTTATA to CTTCTACA by site-directed mutagenesis (primer sequences are listed in Supplementary Table 2). The vector (10 μg) was linearized by *PmeI* and introduced to *P. pastoris* (X-33 strain) by electroporation, for integration into the genome. Expression of the recombinant protein by different transformant colonies was evaluated by western blot, using anti-His antibodies (Sigma, cat # H1029). Recombinant M protein expression in *P. pastoris* was performed as described previously ^49^, with slight modifications. Starter culture was grown overnight at 29 °C at 250 rpm and was used for inoculating 500 mL of BMGY in a 2.5 L baffled flask. The culture was grown at 29 °C at 250 rpm for 72 h. The culture was centrifuged at 3,000 x *g* for 20 min and the BMGY media were discarded. The remaining cell pellet was resuspended in BMMY media (containing 0.5% v/v methanol) to induce protein expression at 15 °C for 20 h. Cells were harvested at 3,000 x *g* for 20 min and the cell pellets were washed with buffer (20 mM Tris pH 7.5, 150 mM NaCl, 5 mM EDTA). Cell pellets were resuspended in lysis buffer (1 mL per 30 g wet weight; 50 mM HEPES pH 8, 150 mM NaCl, 0.25 mM Triton X-100, 5 mM MgCl_2_, 10 % w/v glycerol, 20 mM β-mercaptoethanol, Pierce^TM^ protease inhibitor cocktail tablets, EDTA-free (ThermoFisher Scientific)). Cell pellets were transferred and passed through a 50 mL syringe without a needle into liquid nitrogen. The noodle-shaped pellets were then ground into fine powder in liquid nitrogen, using a mortar and a pestle. Cryoground cells were resuspended in 3 volumes (w/v) of buffer (50 mM HEPES pH 8, 150 mM NaCl, 0.25 mM Triton X-100, 5 mM MgCl_2_, 10% w/v glycerol, 5 mM DTT, SIGMAFAST™ Protease Inhibitor Cocktail Tablets (Merck), 2 mM PMSF). The lysate was centrifuged at 11,899 x *g* for 1.5 h at 4 °C (Avanti Avanti J-26S XP1 centrifuge, JLA-10.500 rotor, Beckman-Coulter). The clarified lysate was loaded onto an equilibrated StrepTactin XT column (IBA Lifesciences) at 4 °C using EP-1 Econo Pump (Bio-Rad). The column was connected to the AKTA Pure FPLC system (Cytiva), and washed with wash buffer (50 mM HEPES pH 8, 150 mM NaCl, 5 mM MgCl_2_, 10% w/v glycerol, 2 mM DTT) until the baseline became stable. Bound proteins were eluted in elution buffer (50 mM HEPES pH 8, 150 mM NaCl, 5 mM MgCl_2_, 10% w/v glycerol, 2 mM DTT, 50 mM biotin) and immediately loaded onto a pre-equilibrated 1 mL cOmplete His-tag purification column (Roche). The column was washed with 12 CVs (column volumes) of wash buffer and the recombinant protein was eluted in buffer containing 250 mM imidazole. The eluted protein fractions (5 mL) were collected and injected onto the 16/600 Superose 6 pg column (Cytiva) equilibrated with SEC buffer (20 mM HEPES pH 7.5, 150 mM NaCl, 5 mM MgCl_2_, 10% w/v glycerol, 2 mM DTT). The total fraction volume of 4 mL from the largest peak was collected (0.25 mg/mL) and snap-frozen in liquid nitrogen for subsequent experiments.

### M resistosome preparation *in vitro*

The purified M protein (1.2 μM) was incubated with AvrM-A (3.6 μM), ATP (1 mM) and either NAD^+^ or 1AD (1.6 mM) in SEC buffer (20 mM HEPES pH 7.5, 150 mM NaCl, 5 mM MgCl_2_, 10% w/v glycerol, 2 mM DTT) at 23 °C for 8 h.

### Negative-stain EM

Six μL of M monomer (130 nM) or M resistosome (the above reaction mixture was diluted in SEC buffer by 10-fold) was applied to a glow-discharged copper grid with formvar/carbon coating (ProSciTech, cat # EMSFCF400-CU-TA), incubated for 1 min at room temperature and blotted with filter paper. The grid was washed with 20 μL of filtered water and blotted with filter paper, and this washing step was repeated once. The grid was then stained with 10 μL of 2% v/v uranyl acetate for 1 min at room temperature and blotted with filter paper. The grid was then air-dried for 1 min. Negative-stain EM images were acquired using a Hitachi HT7700 transmission electron microscope.

### Mass photometry

A microscope glass coverslip was carefully washed with Milli-Q water, ethanol, and isopropanol, dried with clean nitrogen gas and placed onto the instrument with a 6-well gasket attached on top. Protein samples (M monomer or M resistosome) were diluted in SEC buffer (20 mM HEPES pH 7.5, 150 mM NaCl, 5 mM MgCl_2_, 10% w/v glycerol, 2 mM DTT) at room temperature to the stock concentrations of 100-150 nM, immediately before data acquisition on the mass photometer OneMP and analysis using the DiscoverMP software (Refeyn). SEC buffer (15 μL) was added to a well to determine a focal position, followed by addition of a 5 μL protein sample to measure the mass. The mass calibration curve was generated using BSA (bovine serum albumin) monomer (66 kDa), BSA dimer (132 kDa), apoferritin (440 kDa) and thyroglobulin (660 kDa).

### NanoDSF

Purified M protein (1.3 µM) was centrifuged at 10,000 x *g* for 10 min at 4 °C, equilibrated to 20 °C, and loaded into NanoDSF high sensitivity capillaries. The samples were heated in a linear thermal ramp (1 °C/min, 20 °C - 95 °C) with an excitation power of 37 %, using the Prometheus NT.48 (NanoTemper Technologies, Germany). Unfolding transition points were determined from the change in ratio of the emission wavelength of tryptophan and tyrosine fluorescence at 330 and 350 nm. Sample aggregation was determined based on backscattering. Experiments were conducted in triplicate and data were analyzed using the Prometheus PR stability analysis software.

### Cryo-EM sample preparation and data acquisition

Quantifoil Cu 2/1 mesh holey carbon grids were glow-discharged for 60 s at 15 mA by PELCO EasiGlow^TM^ (PELCO). The M resistosome or the M monomer sample was applied on a carbon-coated side of the grid and vitrified using a Thermofisher Vitrobot Mark IV plunge-freezer, with a blotting force of -5, blotting time of 6 s at 5 °C with 90 % humidity. Electron micrographs were acquired using a Titan Krios G2 operating at 300 kV, equipped with Gatan K3 BioQuantum electron detector and a Volta phase plate, with a total electron exposure of 40 e^-^/Å^2^ distributed to 39 frames, and a pixel size of 0.65 Å. Target defocus values were set from -0.6 μm to -1.6 μm for the M resistosome and -1.0 μm to -1.6 μm for the M monomeric protein. Cryo-EM data collection settings are summarized in Table S1.

### Cryo-EM data processing and 3D reconstruction of the M resistosome

Data processing was performed using CryoSPARC (v4.3.1). For 3D reconstruction, raw movies (23,421 movies) were corrected for drift and beam-induced motion, followed by estimation of defocus and CTF-related values. Micrographs with poor CTF-fit resolution, large motion and high astigmatism were removed, leaving 21,152 micrographs for further analysis. Particles were picked automatically using Blob Picker with an expected size ranging from 120 nm to 180 nm in diameter. Particles were extracted and an initial 2D classification was performed (n = 50). Two 2D classes were selected as templates for re-picking particles by Template Picker, and extracted particles were subjected to two further rounds of 2D classification (n = 50 and n = 100) to remove bad particles, yielding a subset of 193,972 particles. After *ab-initio* reconstruction (n = 4), this subset of particles was further classified using heterogeneous refinement (n = 4). Particles from the best 3D class (181,574 particles) were used for 3D refinement with C_2_ symmetry, as per reconstruction of the RPP1 resistosome ^15^. After CTF refinement and post-processing, the resolution of the final cryo-EM map was 2.23 Å. Because the EM density corresponding to the M^LRR^:AvrM-A region was poorly resolved, we carried out further processing around this region. C_4_ symmetry was applied to refine the particles, followed by C_4_ symmetry expansion of these particles. 3D classification was then performed with a focused mask (generated by UCSF ChimeraX v1.8) around the target region. A subset of 124,664 particles was selected for local 3D refinement with C_1_ symmetry, resulting in reconstruction of the M^LRR^:AvrM-A region with 2.71 Å resolution. Both maps were sharpened using DeepEMhancer ^64^. Data processing workflow is shown in Figure S3.

### Model building and refinement of the M resistosome

Initial models for M^TIR^, M^NB-ARC^ and M^LRR^ were obtained using SWISS-MODEL ^65^, with the L6^TIR^ crystal structure (PDB: 6OIZ), the RPP1 cryo-EM structure (PDB: 7LJV), and the L6^LRR^ AlphaFold DB model as templates, respectively. These models were docked into the consensus cryo-EM density map using UCSF ChimeraX ^66^. The model was built to the beginning of M^LRR^ (up to residue 624). Then, the final EM map (the composite EM map) was generated by combining the consensus EM map and the local EM map of the M^LRR^:AvrM-A region, using vop commands in UCSF ChimeraX. This composite EM map was used to model the AvrM-A and the rest of the M^LRR^ region, where the AvrM-A crystal structure (PDB: 4BJN) was docked into the density in UCSF ChimeraX. Terminal M^LRR^ residues were manually built using COOT, using the M^LRR^ models generated by SWISS-MODEL, AlphaFold 2 and AlphaFold 3 as guides. The two models (M^TIR:NB-ARC^ and M^LRR:AvrM-A^) were combined to generate the final M resistosome model. The final model was refined against the composite EM map using ISOLDE ^67^ in UCSF Chimera X, followed by real-space refinement in Phenix ^68^.

### Cryo-EM data processing and modelling of the M monomer

Data processing was performed using CryoSPARC (v4.6.0). Raw movies (23,125 movies) were obtained and corrected for drift and beam-induced motion, followed by estimation of defocus and CTF-related values. Micrographs with low CTF resolution, large motion and strong astigmatism were filtered out, leaving 22,194 micrographs for subsequent processing. The micrographs were de-noised for particle picking using Micrograph Denoiser. Particles were manually picked using a subset of 195 micrographs, followed by 2D classification (n = 10). Four 2D classes were selected as templates for Template Picker using a subset of 1,000 micrographs and two rounds of 2D classification were performed (n = 50 and n = 7). Finally, three of the 2D classes were selected as templates for further picking of particles from all the micrographs. After seven rounds of 2D classification, a subset of five 2D classes containing 113,296 particles was selected. This subset of particles was used for *ab-initio* reconstruction (n = 4), followed by heterogeneous refinement (n = 5) using the best volume from the *ab-initio* reconstruction job and four decoy volumes. This process was repeated once to generate an initial model with interpretable features. This initial model, along with four decoy volumes, was then used for heterogeneous refinement with a subset of particles (384,947 particles) that was obtained after the sixth round of 2D classification, in order to enrich for well-defined particles. From this heterogeneous refinement, 124,886 particles from the best volume were used to generate three models by *ab-initio* reconstruction (n = 3), resulting in a volume with 55,512 particles that were selected for re-extraction at 320 pixels without binning, followed by final *ab-initio* reconstruction (n = 1) and non-uniform refinement (C_1_). The individual domains of M (NBD, HD1, WHD and LRR from a protomer of the M resistosome) were docked into the cryo-EM map of the M monomer and the model was refined using Phenix real-space refinement.

### Expression and purification of effector proteins

With the aim of identifying the residues important for controlling M recognition and taking into consideration all of the data previously reported ^32,35,69^, the *AvrM-A* and *avrM* genes were truncated and engineered to encode proteins from residues 103-343 and 46-280, respectively. The gene fragments of *avrM* and *AvrM-A* and their mutant variants were cloned into the pMCSG7 expression vector. A 6xHis-tag was engineered onto the N-termini of the effector proteins, to facilitate purification by nickel affinity chromatography. All expression vectors were verified by Sanger sequencing. Protein expression and purification were performed as previously described ^35^. In brief, proteins of interest were purified utilising nickel affinity chromatography and SEC. The purity of the proteins was estimated by Coomassie Blue-stained SDS-PAGE.

### SEC-MALS (size-exclusion chromatography coupled with multi-angle light scattering)

SEC-MALS was performed using an in-line Superdex 200 Increase 5/150 GL column (Cytiva) coupled with a DAWN HELEOS II 18-angle light-scattering detector combined with an Optilab TrEX refractive index detector (Wyatt Technology) and connected to a Prominence UFLC (Shimadzu). Purified protein at 1.0 mg/mL was injected (40 μL) and separated at a flow rate of 0.25 mL/min in SEC buffer (10 mM HEPES pH 7.5, 150 mM NaCl, 1.0 mM DTT) at 22 ℃, with data collected at 0.2 s intervals. The ASTRA6.1 software (Wyatt Technology) was used for the calculations of the molecular mass across the protein elution peak. A constant value of 0.185 was set for the refractive increment (dn/dc values) in the molecular mass calculations, because dn/dc was likely to be unchanged for non-modified proteins ^70^.

### NMR analysis of NAD^+^-cleavage

Full-length M and L6 proteins were expressed and purified as described above, except that the final SEC buffer excluded 10 % w/v glycerol. M (residues 21-1305; 0.8 µM) or L6 (residues 29-1294; 1.5 µM), were incubated either with 2 mM NAD^+^ in the presence or absence of 2 mM ATP, and both conditions were repeated with 8 µM of their respective effectors AvrM-A (residues 103-343) or AvrL567-A (residues 23-150) in SEC buffer (10 mM HEPES pH 7.5, 150 mM NaCl). Reactions were quenched at 0 h and 24 h at room temperature. To quench the reactions, 25 µL of the reaction mixture were added to 175 µL NMR quenching buffer (1% HCl, 20 µM sodium trimethylsilylpropanesulfonate, 5% v/v deuterium oxide and 50 µL SEC buffer). To remove any precipitant, the samples were centrifuged at 10,000 x *g* for 10 min and the supernatant was transferred into 3 mm Bruker SampleJet tubes (Bruker). Spectra were recorded using an Avance Neo 900 MHz NMR spectrometer (Bruker) equipped with a cryogenically cooled probe. Data was collected with water suppression through excitation sculpting (180° water-selective) and a flip back pulse (zgesfpgp). The spectra were acquired using 512 scans with an interscan delay of 1 s, at 298 K. All spectra were processed and analyzed using TopSpin™ (Bruker). NAD^+^ cleavage was validated by detecting nicotinamide (Nam). No quantification was performed as NAD^+^ was either completely consumed or unaffected (validated by no Nam build up). All spectra were referenced to DSS (sodium trimethylsilylpropanesulfonate). For TIR-derived product determination, reactions were compared to standards of cADPR, 2’cADPR, 3’cADPR, pRib-AMP, ADPR, ATP, NAD^+^, or Nam at 50-75 µM in reaction buffer.

### MALDI-TOF MS analysis of NAD^+^-cleavage products

From the NADase reactions above, 5 µL was taken from the 24 h time-points and added to 5 µL ddH_2_O to decrease NaCl concentration. To remove the protein and quench the reaction, samples were boiled at 90 °C for 5 min, centrifuged at 10 000 x *g* for 10 min and the supernatant was transferred to a new tube. α-Cyano-4-hydroxycinnamic acid (CHCA) in acetonitrile was diluted by ten-fold in ethanol: acetone: 0.1% v/v trifluoroacetic acid (TFA) (6:3:1) solution and used as a matrix. 2 µL matrix was mixed with 1 µL sample. Each sample was spotted twice on a MTP 384 ground steel MALDI plate. MS was performed on a Bruker TIMS TOF Flex - MALDI-2 (Bruker) with TIMS TOF Control 5.1; a calibration was performed using ESI (electrospray ionization) calibration solution (Fluka 63606), and the error was less than 0.5 ppm for all samples. Laser power was set to 62% at a repetition rate of 1000 Hz. An MS scan was performed in positive QqTOF mode over a range of 200 - 2000 m/z. MS parameters were tuned to detect compounds from 200 – 1500 m/z using funnel 1 RF: 350 Vpp; isCID: 0.0 eV; funnel 2 RF: 350 Vpp; multipole RF: 350 Vpp; collision energy: 10 eV; collision RF: 1500 Vpp; quadrupole settings: ion energy: 5 eV; low mass: 250 m/z; focus pre-TOF settings: transfer time: 80 µs; pre-pulse storage: 10 µs; and detection was set to focus mode. MS/MS fragmentation was performed with a width range 0.8 m/z and 15 eV collision energy, and summed with second round collisions at 35 eV.

### Fluorescence-based NADase assays

A fluorescent NAD^+^ analogue, 1,N^6^-ethenoNAD (εNAD), was used as a substrate. An increase in fluorescence from 1,N^6^-ethenoadenine diphosphate ribose occurs upon cleavage of the quenching nicotinamide group. The stocks of εNAD were prepared in buffer (20 mM HEPES pH 7.5, 150 mM NaCl, 5 mM MgCl_2_, 2 mM DTT). Each protein sample was diluted to a final concentration of 300 nM. εNAD (0.1 mM) was added into the reaction mix in each of a 96-well plate containing protein samples (the 1:3 molar ratio of NLR:effector proteins) and ATP (1 mM). Fluorescence intensity was measured using an excitation wavelength of 310-330 nm and emission wavelength of 390-410 nm by a CLARIOstar microplate reader. Readings were recorded every 1-3 min for 12 h at 25 °C. The change in fluorescence intensity over time was calculated based on slopes of the linear component of the curves.

### Vectors and constructs for *in planta* analysis of AvrM and avrM effector variants on M activity

AvrM (residues 108-343) and avrM (residues 46-280) sequences were cloned into the pEarleygate 201 vector ^71^ with a cauliflower mosaic virus (CaMV) 35S promoter. This cloning also engineered a sequence encoding an N-terminal hemagglutinin (HA) epitope tag for immunoblot detection of effector proteins. Site-directed mutagenesis PCR was performed to engineer the combined mutations in *avrM* and *AvrM-A* genes contained within the pEarleygate 201 vector. Mismatch primers were designed to introduce the desired mutation/s in the effector genes. The mutant genes were prepared and electroporated in *E. coli* (DH10B) and the integrity of the resultant constructs was confirmed by Sanger sequencing.

### Transient expression and immunoblot analysis of AvrM-A, avrM and variants

Transgenic tobacco (*N. tabacum* cv. White Burley) containing the *M* gene was grown in a growth chamber (12 h light/12 h dark at 23 °C). *Agrobacterium tumefaciens* (strain GV3101 pMP90) cells containing the *AvrM-A* or *avrM* gene constructs were grown in LB supplemented with 25 mg/mL rifampicin, 25 mg/mL gentamycin and 50 mg/mL kanamycin at 28 °C for 2 days. Cells were pelleted and resuspended in infiltration media (10 mM MgCl_2_, 10 mM MES pH 5.6, 200 µM acetosyringone), to give a final OD_600_ = 1.0, and incubated at room temperature for 2-3 h. Infiltration of rapidly expanding tobacco leaves from 6-week-old plants were carried out with a blunt needless syringe. Plants were then returned to the growth chamber for 2-4 days. For immunoblot analysis, to check for protein accumulation and stability *in-planta*, we used *N. benthamiana*, as other confounding immuno-reactive peptides were detected in *N. tabacum* leaves at the same molecular weight as AvrM-A, avrM and variants. Leaf tissue (∼ 1 cm diameter discs) was collected from the middle of an infiltrated sector 48 h after agroinfiltration and stored at -80 °C. Tissue was ground up in 100 µL of 2x Laemmli extraction buffer, centrifuged at 23,000 x *g* for 10 min at 4 °C and the supernatant fraction collected and heated to 95 °C prior to SDS-PAGE. Proteins were separated by SDS-PAGE and transferred to nitrocellulose membranes (Pall). Membranes were blocked in 5% w/v skim milk and probed with anti-HA antibody (Jomar Life Research), followed by goat anti-mouse antibodies conjugated with horseradish peroxidase (Thermofisher Scientific). Antibody binding was detected with a SuperSignal West Pico chemiluminescence kit (Thermofisher Scientific) following the manufacturer’s instructions. Membranes were stained with Ponceau-S to assess for equivalent protein loading.

### Yeast two-hybrid assays

*M* (residues 21-1305) was cloned into pGBT9 and pGADT7 vectors (Clontech), as described previously ^32^. *AvrM-A* (residues 108-343), *avrM* (residues 46-280) and the corresponding mutants were cloned into pGBT9 and pGADT7 vectors (Clontech) as *EcoRI-BglII* fragments. All constructs were verified by sequencing. Yeast (HF7c strain) transformation and growth assays were performed as described in the Yeast Protocols Handbook (Clontech). Yeast protein extraction for immunoblot analysis was performed following a post-alkaline extraction method as described previously ^72^. Protein fusion detection was performed using anti-HA-HRP (Roche, clone 3F10), and anti-GAL4 DNA BD (SIGMA, G3042) antibodies.

### Transient expression in *N. benthamiana* for cell-death assessment

The pAM-PAT-35S-GWY-YFP binary vector, harboring AvrM-A without a signal peptide, and the pEarleyGate 100 binary vector, carrying a genomic *M* fragment fused in-frame with an HA epitope tag-coding sequence, used for the effector-dependent cell-death analysis, were described previously ^49,73^. *DpnI*-mediated site-directed mutagenesis, using Phusion High-Fidelity DNA polymerase (M0530L; NEB), was performed on the M construct. To test autoactivity of the TIR domains, M (residues 1-263) and the related M1 (residues 1-248) sequences were amplified for Gateway cloning BP reaction (TheroFisher Scientific) to clone into pDONR207. The fragments were then subcloned into binary vectors, pAM-PAT-35S-GWY-YFP and modified pAM-PAT-35S-GWY-Myc carrying the hSARM1^tSAM^ or hSARM1^tSAM(5Mut)^ fusions, as described previously ^14^. The M1(D540V)-YFP binary construct ^73^ was used as a positive control. *Agrobacterium* strain GV3101 pMP90 and GV3103 pMp90 RK were used to transform the binary constructs described above. 4-week-old plants grown at 23 ℃ with a 16 h/8 h light/dark photoperiod were used for transient expression. Transformed *Agrobacterium* was cultured at 28 ℃ with shaking overnight in LB liquid media supplemented with 25 mg/mL rifampicin, 15 mg/mL gentamicin and either 50 mg/mL kanamycin or 25 mg/mL carbenicillin. The cells were harvested and resuspended with infiltration buffer (10 mM MES pH 5.6, 10 mM MgCl_2_, 150 mM acetosyringone). The cells were incubated for 2 h at room temperature and diluted to a final concentration at OD_600_ = 1.0 for infiltration. Leaf samples were taken 24 h after infiltration for protein extraction to perform western blots. Leaf samples for cell death evaluation were taken 5 days after infiltration.

Total protein from plant leaf tissues was extracted for immunoblot analysis using extraction buffer (125 mM Tris-HCl pH 7.5, 4% w/v SDS, 20% w/v glycerol, 0.2 M DTT, 0.02% w/v Bromophenol Blue). Proteins were separated by SDS-PAGE and transferred to nitrocellulose membranes. Membranes were blocked by 5% w/v skimmed milk, followed by overnight incubation with anti-HA-Peroxidase, High Affinity (Roche, ref. 12013819001), anti-GFP, or anti-myc antibodies. Ponceau-S was used to stain membranes and antibody signals were detected using the Super Signal West Femto chemiluminescence kit (ThermoFisher Scientific).

---

**Figure S1.**
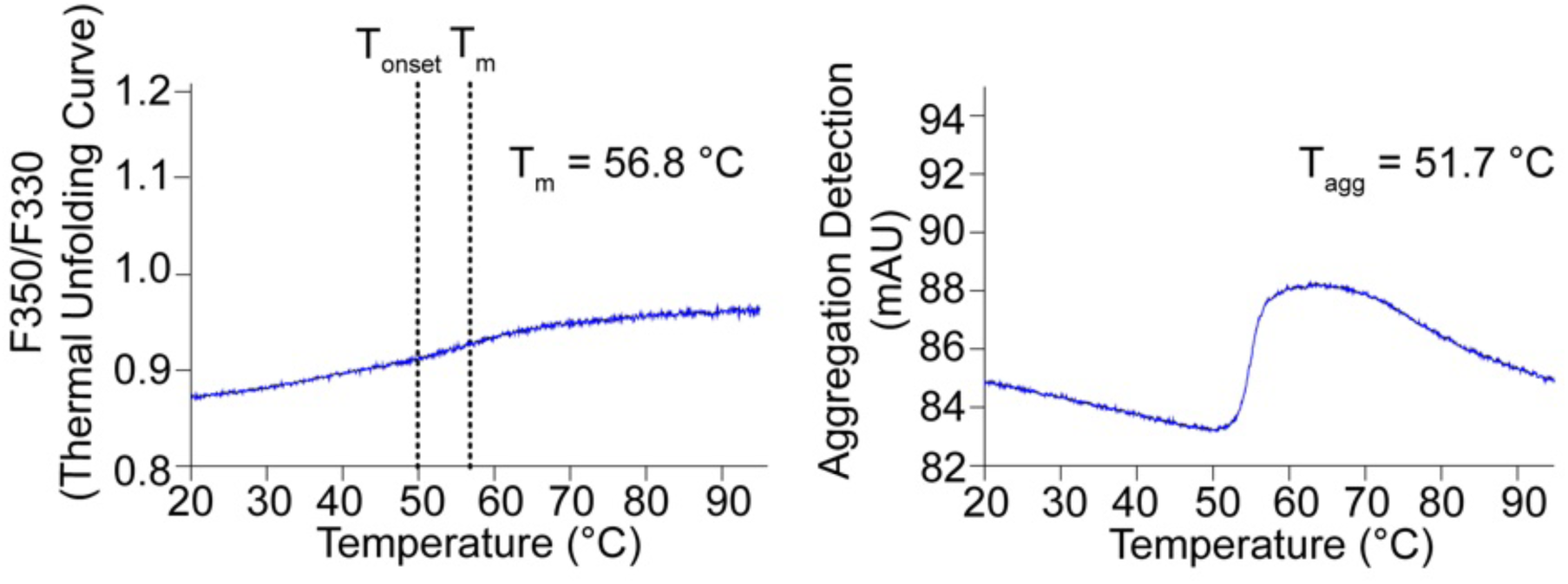
Biophysical characterization of the flax M protein. M protein at 1.3 µM was analyzed by NanoDSF. The melting temperature (T_m_) and temperature at which the protein unfolds (T_onset_) (left; indicated by vertical lines), and the aggregation temperature (T_agg_) (right), are indicated.

**Figure S2.**
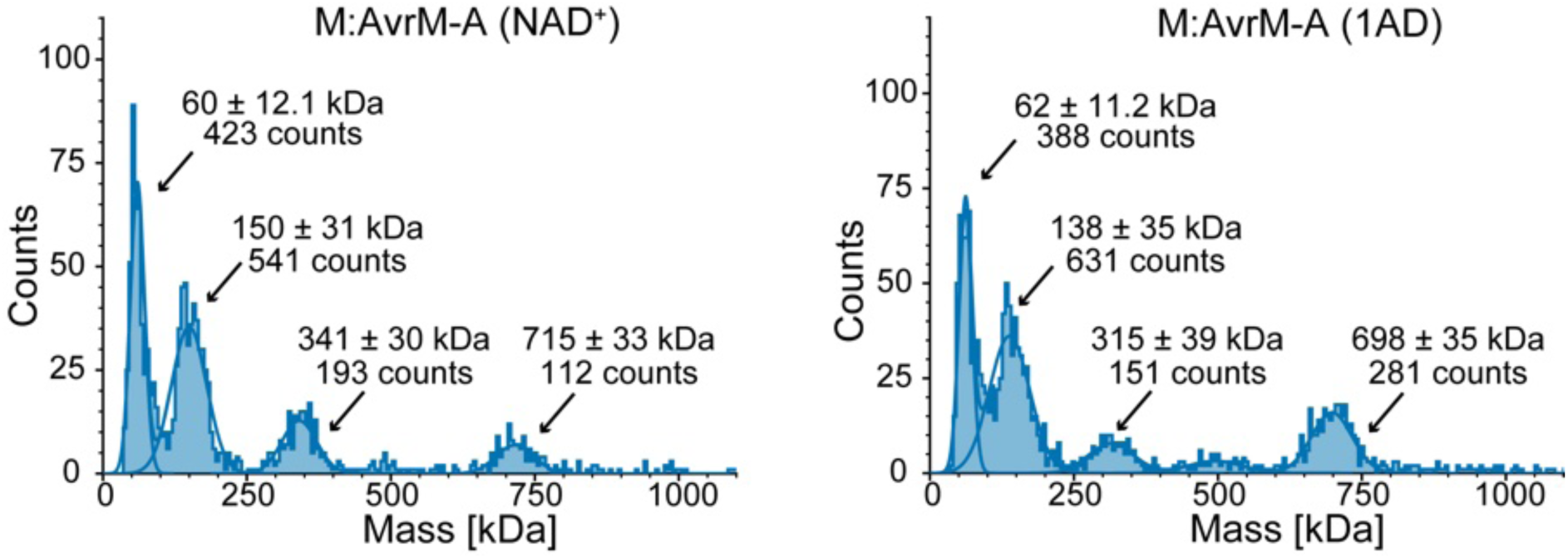
Effects of AvrM-A, ATP and NAD^+^ analogues on the M resistosome formation, assessed by mass photometry. Effector, ATP, and NAD^+^ or 1AD were incubated with the full-length M protein *in vitro* at 23 °C for 8 h. The proteins were diluted to 25 - 37.5 nM in SEC buffer (20 mM HEPES pH 7.5, 150 mM NaCl, 5 mM MgCl_2_, 10% w/v glycerol, 2 mM DTT) during recording on the mass photometer OneMP (Refeyn) and analyzed using the DiscoverMP software.

**Figure S3.**
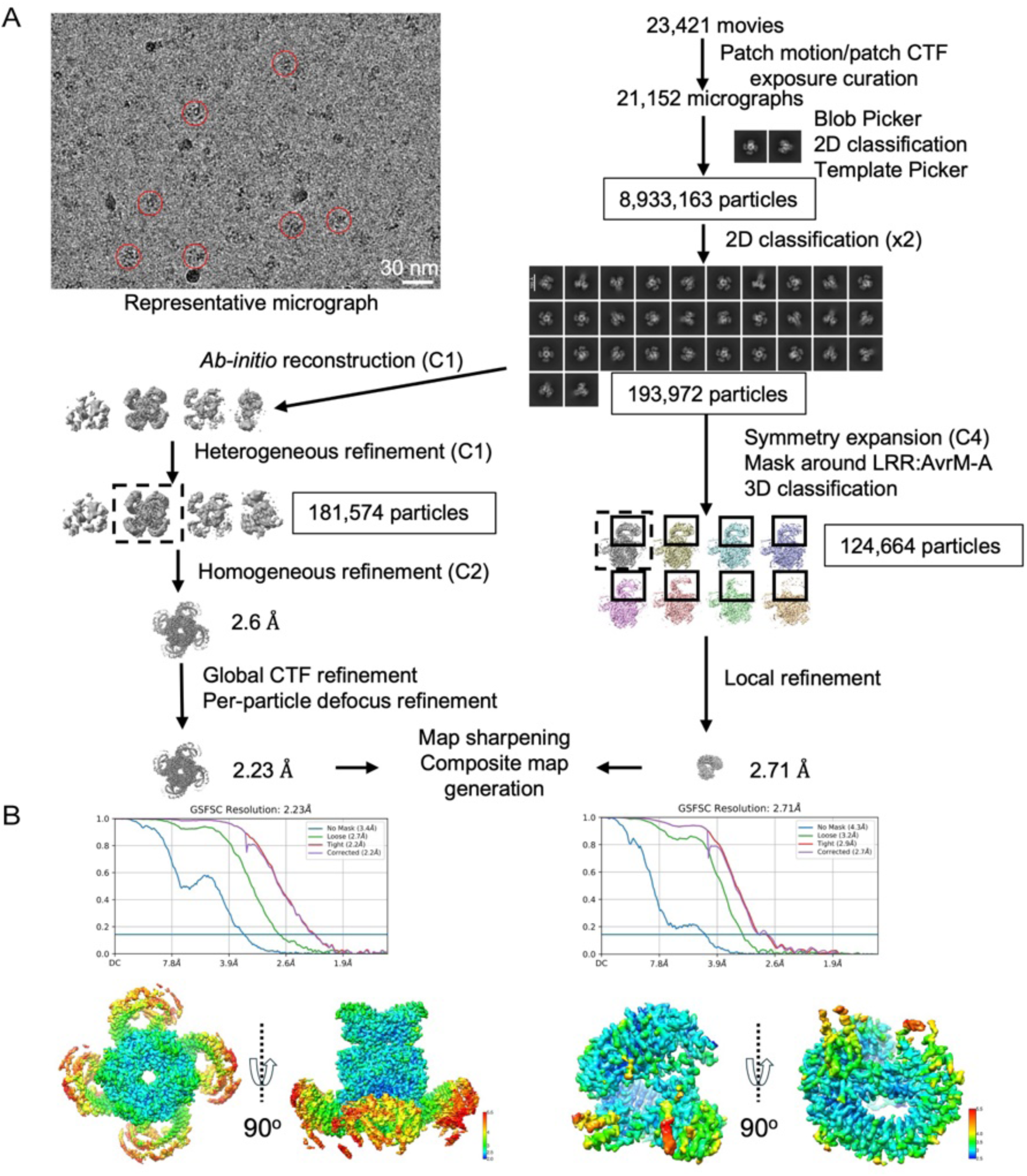
Cryo-EM reconstruction of the M resistosome in complex with the AvrM-A protein. **(A)** Data processing workflow for the M:AvrM-A complex structure determination. A representative cryo-EM micrograph of the M:AvrM-A resistosome is shown on top left, with particles marked in red circles. Scale bar = 30 nm. **(B)** GSFSC (gold-standard Fourier shell correlation) at 0.143 and an estimated local resolution map for the final reconstruction of the M resistosome (left) and the M^LRR^:AvrM-A region (right).

**Figure S4.**
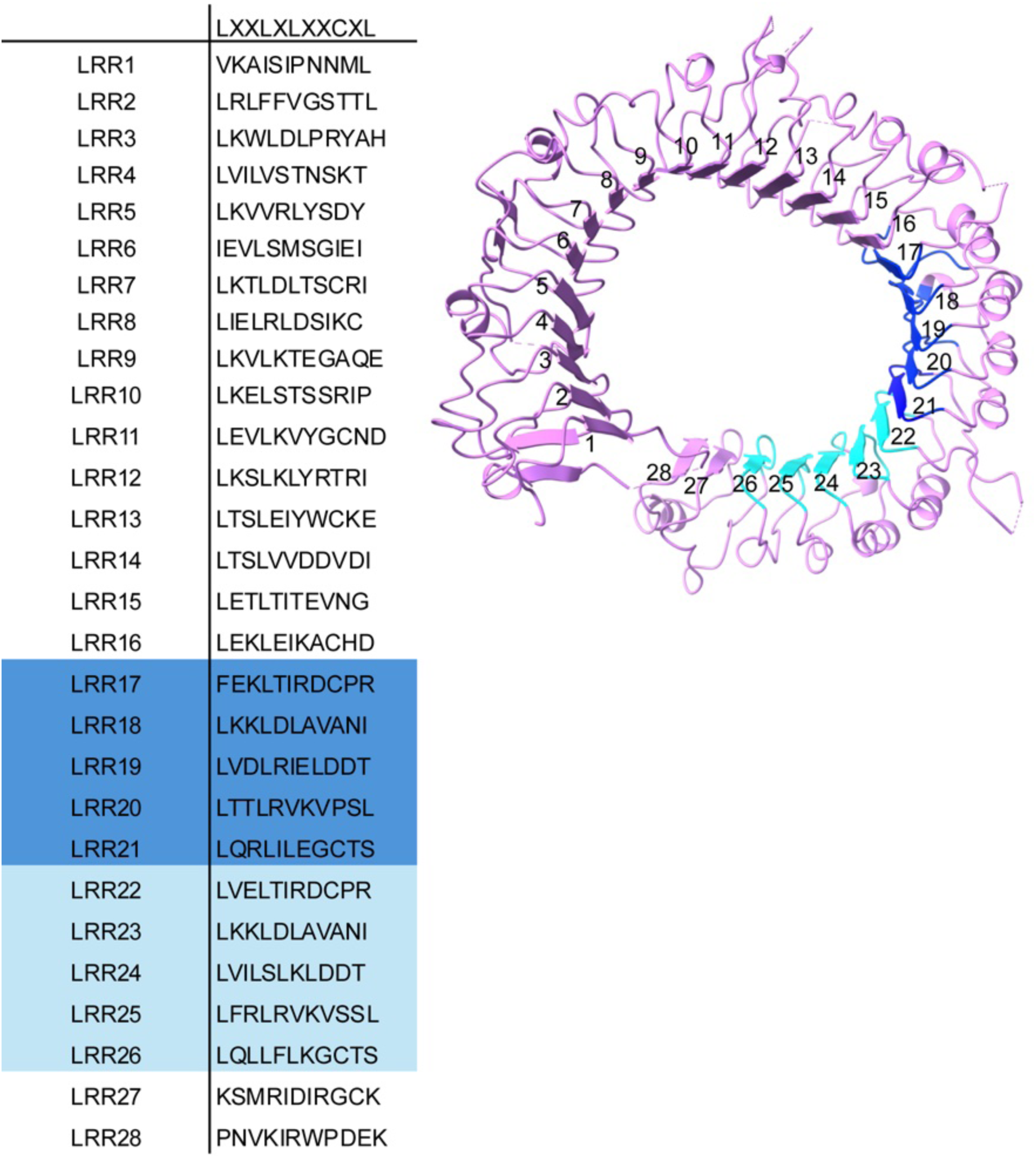
M LRR motif sequences. The table lists every LRR motif sequence for M. The consensus LRR sequence is LxxLxLxx(N/C)xL ^74^. The flax M TNL contains 28 LRRs. At the C-terminus of the LRR region, there are two copies of a tandem repeat (TR) sequences, with TR1 (LRR17-LRR21; highlighted in blue) involving 147 amino acids (residues 980-1126) and TR2 (LRR22-LRR26; highlighted in cyan) involving 149 amino acids (residues 1127-1275). These TR sequences share 77 % amino acid sequence identity and probably arose as a result of unequal intragenic recombination (90 % nucleotide sequence identity).

**Figure S5.**
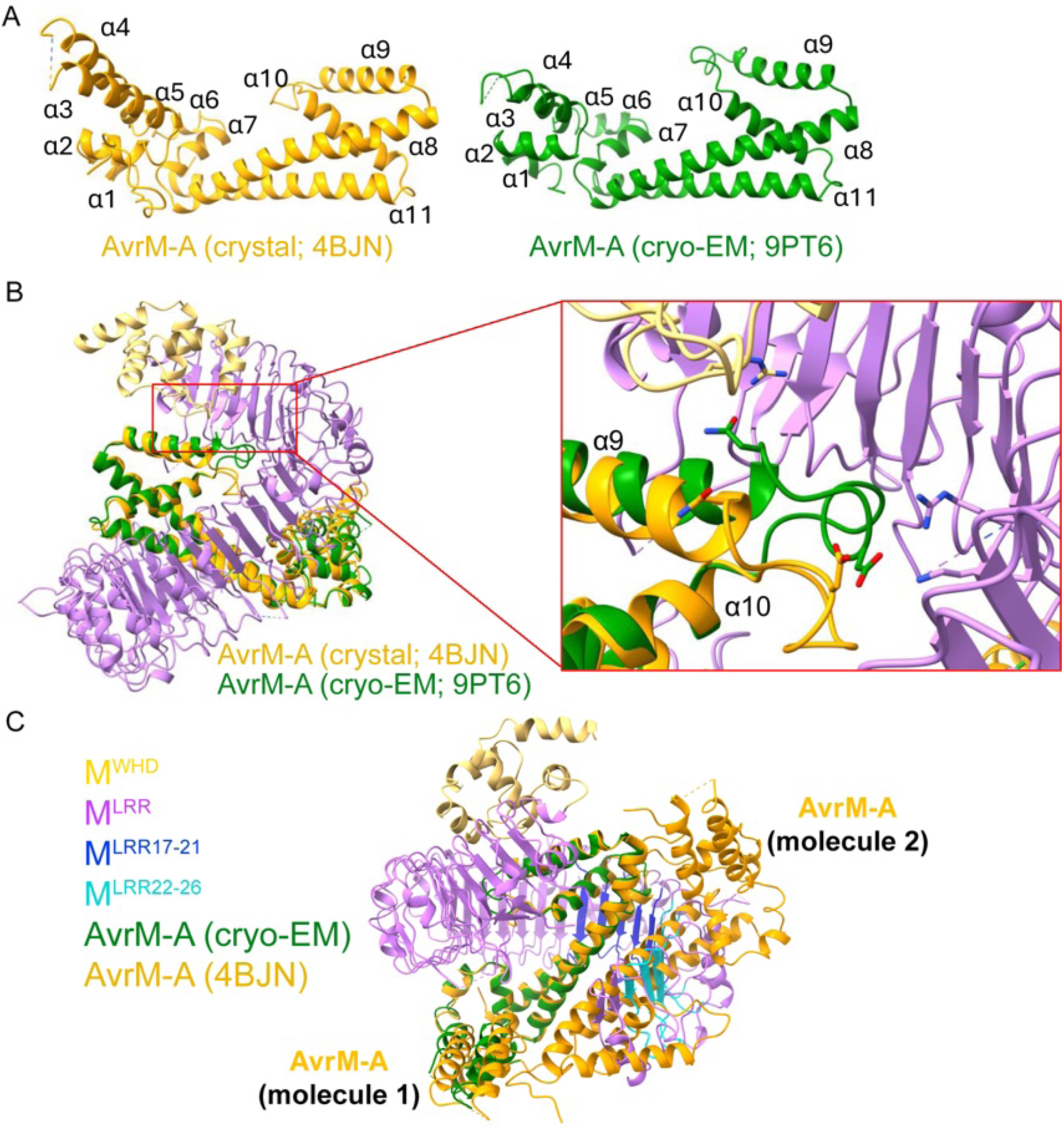
Structural characterization of the WHD and LRR domains of the M protomer interacting with the AvrM-A effector protein. **(A)** Crystal structure of the AvrM-A protein (left; PDB: 4BJN, chain A) and cryo-EM structure of the AvrM-A protein from the M resistosome (right, chain E), shown in cartoon representation. The elements of secondary structure are labeled. **(B)** Superposition of the AvrM-A protomer crystal structure (orange) on the cryo-EM AvrM-A structure (green). RMSD (root mean square deviation) is 0.981 Å for 124 Cα atoms. Overall view (left) and a close-up view (right) are shown. Compared to the crystal structure, the α9 helix and the α9-α10 loop of AvrM-A in the M resistosome are positioned differently (closer towards M^WHD^ and M^LRR^ in the M resistosome). The different domains of the M resistosome are colored: M^WHD^, yellow; M^LRR^, magenta; and AvrM-A, green. **(C)** Superposition of the cryo-EM structure of the M:AvrM-A protein complex with the crystal structure of the AvrM-A protein dimer (orange; PDB: 4BJN). The different domains of the M resistosome are colored: M^WHD^, yellow; M^LRR^, magenta; M^LRR^ TR1, blue; M^LRR^ TR2, cyan; and AvrM-A, green. The non-interacting AvrM-A protein molecule in the dimer (molecule 2) would have steric clashes with M^LRR^ TR2.

**Figure S6.**
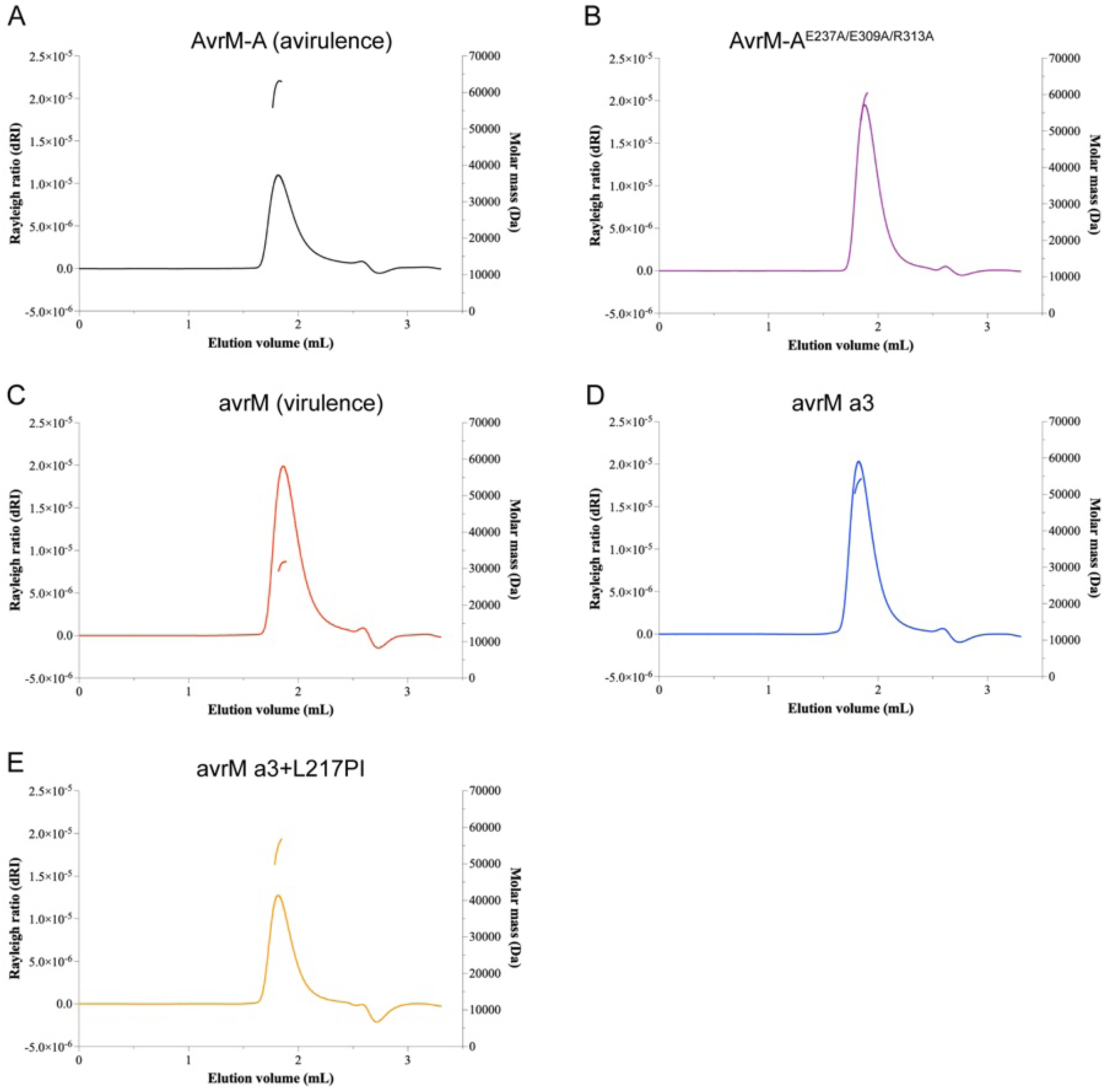
SEC-MALS analysis of AvrM-A and wild-type and mutant avrM proteins. Samples tested include **(A)** wild-type (avirulence) AvrM-A; **(B)** AvrM-A (E237A/E309A/R313A); **(C)** wild-type (virulence) avrM; **(D)** avrM a3; and **(E)** avrM a3+L217PI. The avrM a3 mutant contains three mutations (R170K/S179L/T247I) in the α8 and α11 helices that mimic AvrM-A. The avrM a3+L217PI mutant contains the additional L217I mutation and an extra proline residue, to resemble AvrM-A in the α9-α10 loop region. The protein variants at 10-20 mg/mL eluted in a peak of a molar mass consistent with a dimeric form (∼58 kDa), except for avrM, which eluted in a peak of molar mass indicating a monomeric form (∼29 kDa).

**Figure S7.**
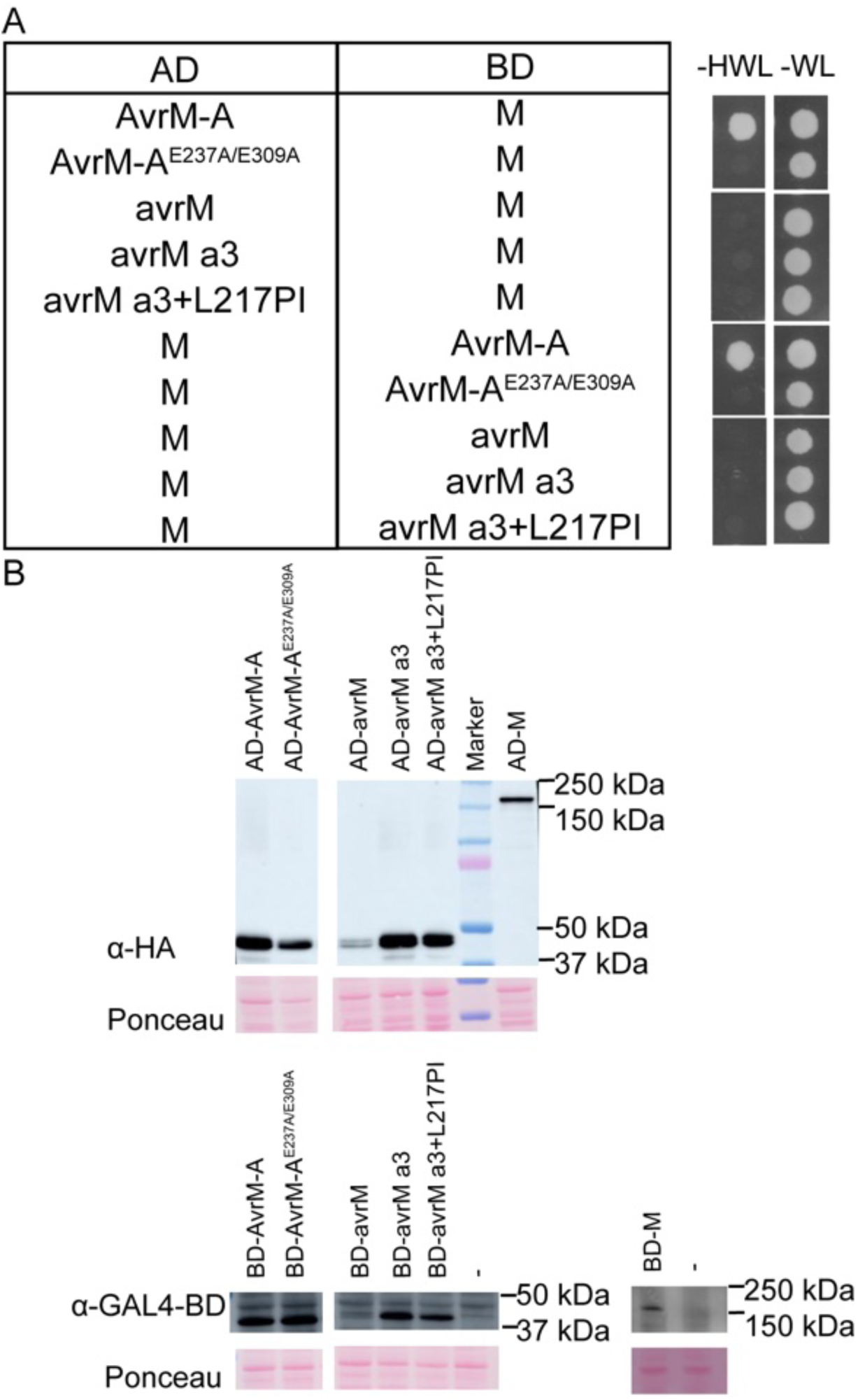
Physical interaction of M with AvrM and avrM mutants in yeast two-hybrid assays. Samples tested include: wild-type (avirulence) AvrM-A; AvrM-A (E237A/E309A); wild type (virulence) avrM; avrM a3; and avrM a3+L217PI. The avrM a3 mutant contains three mutations (R170K/S179L/T247I) that mimic AvrM-A in the α8 and α11 helices. The avrM a3+L217PI mutant contains the additional L217I mutation and an extra proline residue, to resemble AvrM-A in the α9-α10 loop region. **(A)** Growth of yeast cells co-expressing GAL4-AD and GAL4-BD fusions of M and AvrM or avrM mutants (and vice versa) on non-selective media lacking tryptophan and leucine (-WL); or selective media additionally lacking histidine (-HWL). Growth on -HWL represents positive interaction. **(B)** Immunoblot detection of GAL4-AD and GAL4-BD fusions of M, AvrM, avrM and mutants. Proteins were detected using anti-HA (GAL4-AD fusions) and anti-GAL4 DNA BD (GAL4-BD fusions) antibodies. Negative control is indicated by a dash and corresponds to untransformed yeast. Protein loading is indicated by red Ponceau staining.

**Figure S8.**
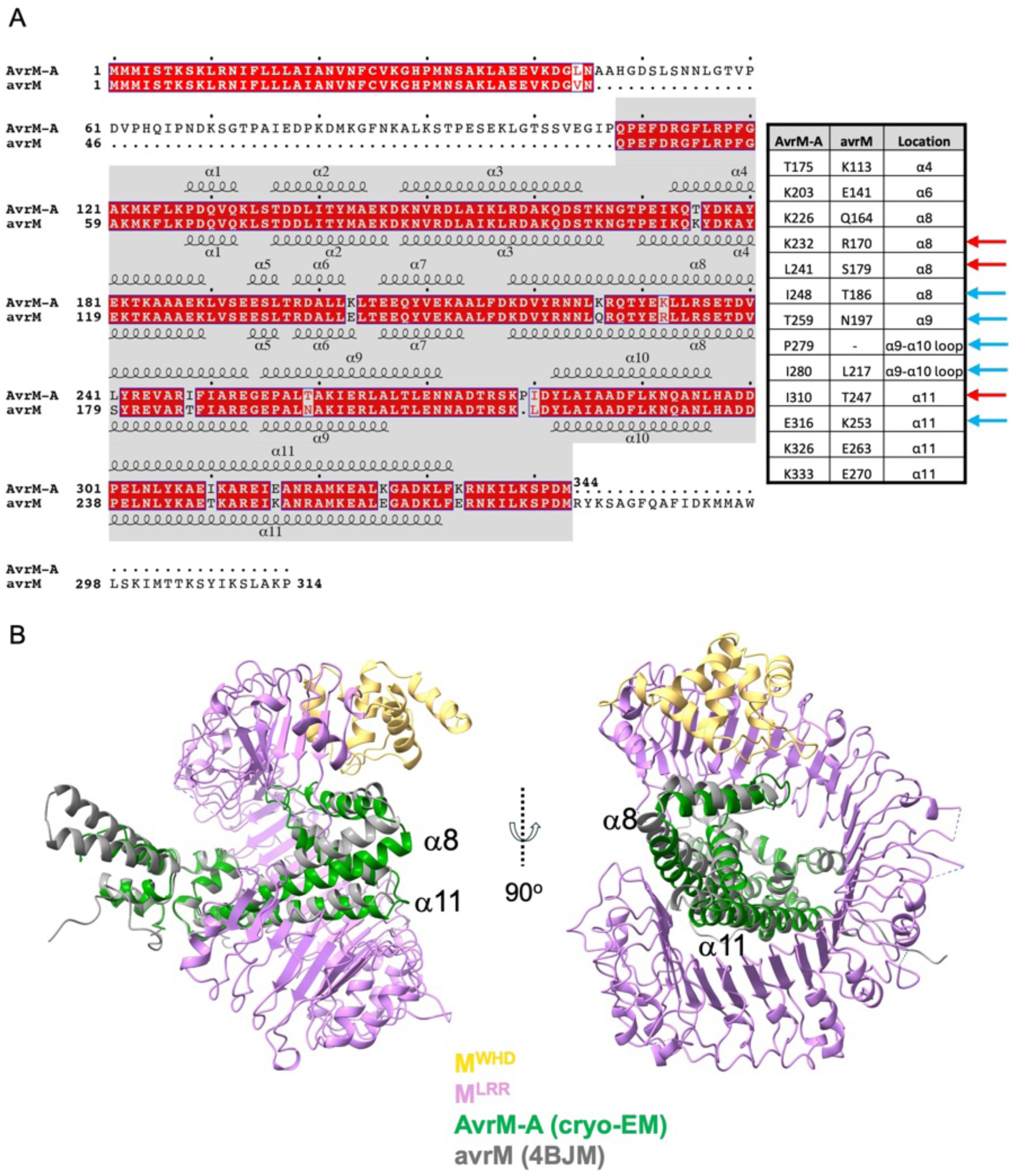
Sequence and structural comparison of AvrM-A and avrM. **(A)** Alignment of protein sequences between AvrM-A (residues 1-343) and avrM (residues 1-314) from the flax rust fungus *M. lini*. 13 polymorphic residues are summarized in the table (right) with a description of their location. In the table, red arrows indicate the three residues used to generate the avrM a3 mutant and cyan arrows indicate residues that were further added to the avrM a3 mutant. **(B)** Superposition of the cryo-EM structure of the M:AvrM-A protein complex with the crystal structure of the avrM-A protein (grey; PDB: 4BJM; chain A). The different domains of the M resistosome are colored: M^WHD^, yellow; M^LRR^, magenta; and AvrM-A, green.

**Figure S9.**
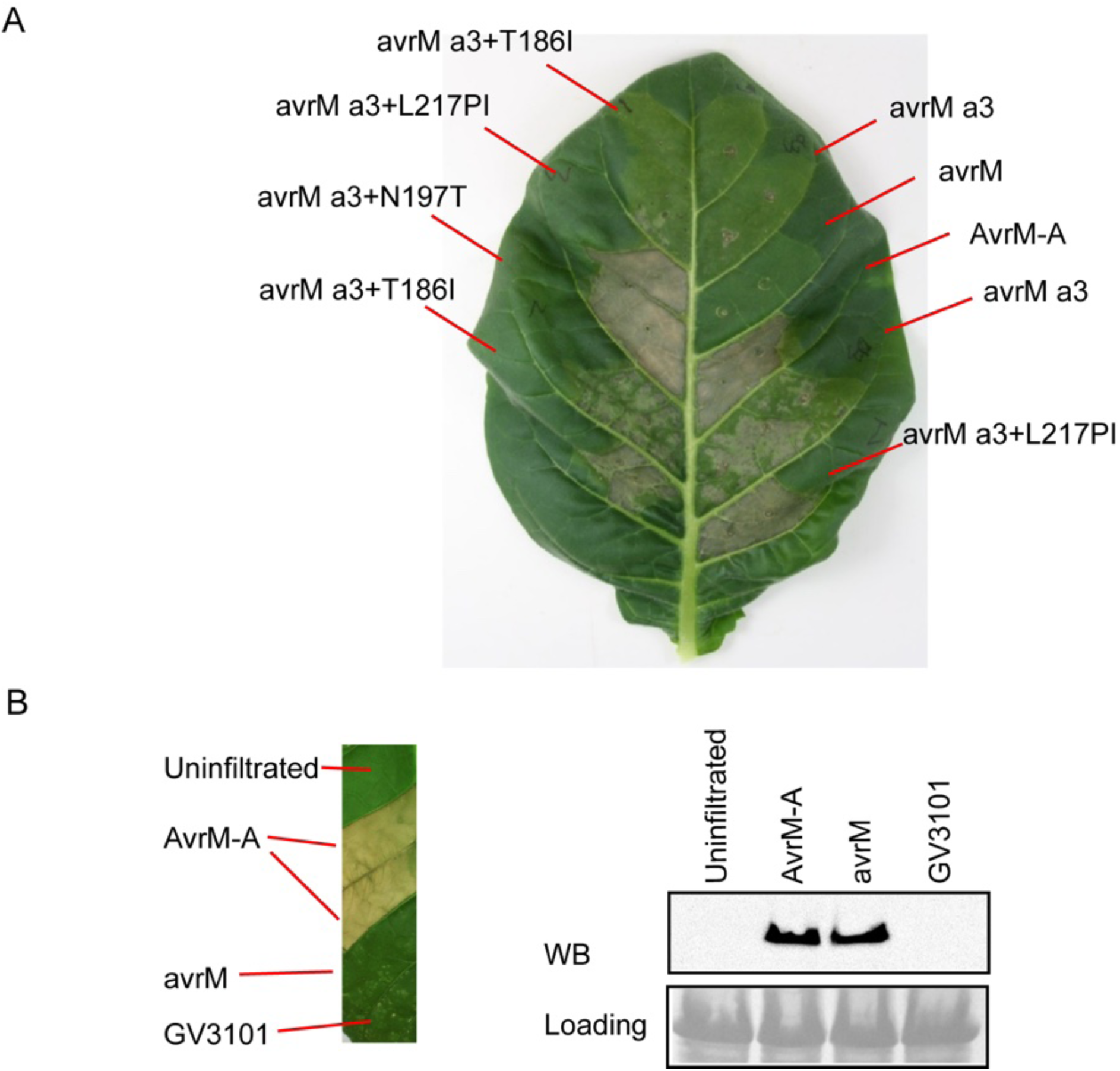
Effects of transient expression of avrM (virulence) effector variants on M-dependent cell death in leaves of *M*-transgenic *N. tabacum*. **(A)** Similar to Figure 4E, effectors tested include wild-type avirulence AvrM-A (positive control), virulence avrM (negative control), and mutant avrM variants carrying a3 (R170K/S179L/T247I), and avrM a3+ variants (a3+K186I, a3+N197T or a3+L217PI). It is noted that the avrM a3 mutant could induce M-dependent HR in some, but not all infiltration experiments. **(B)** Leaf infiltration experiments with control samples: uninfiltrated, AvrM-A, avrM and GV3101 pMP90 (untransformed *Agrobacterium* strain infiltrated as a negative control). Protein expression was validated by western blotting (WB). Equivalent sample loading is shown by SDS-PAGE.

**Figure S10.**
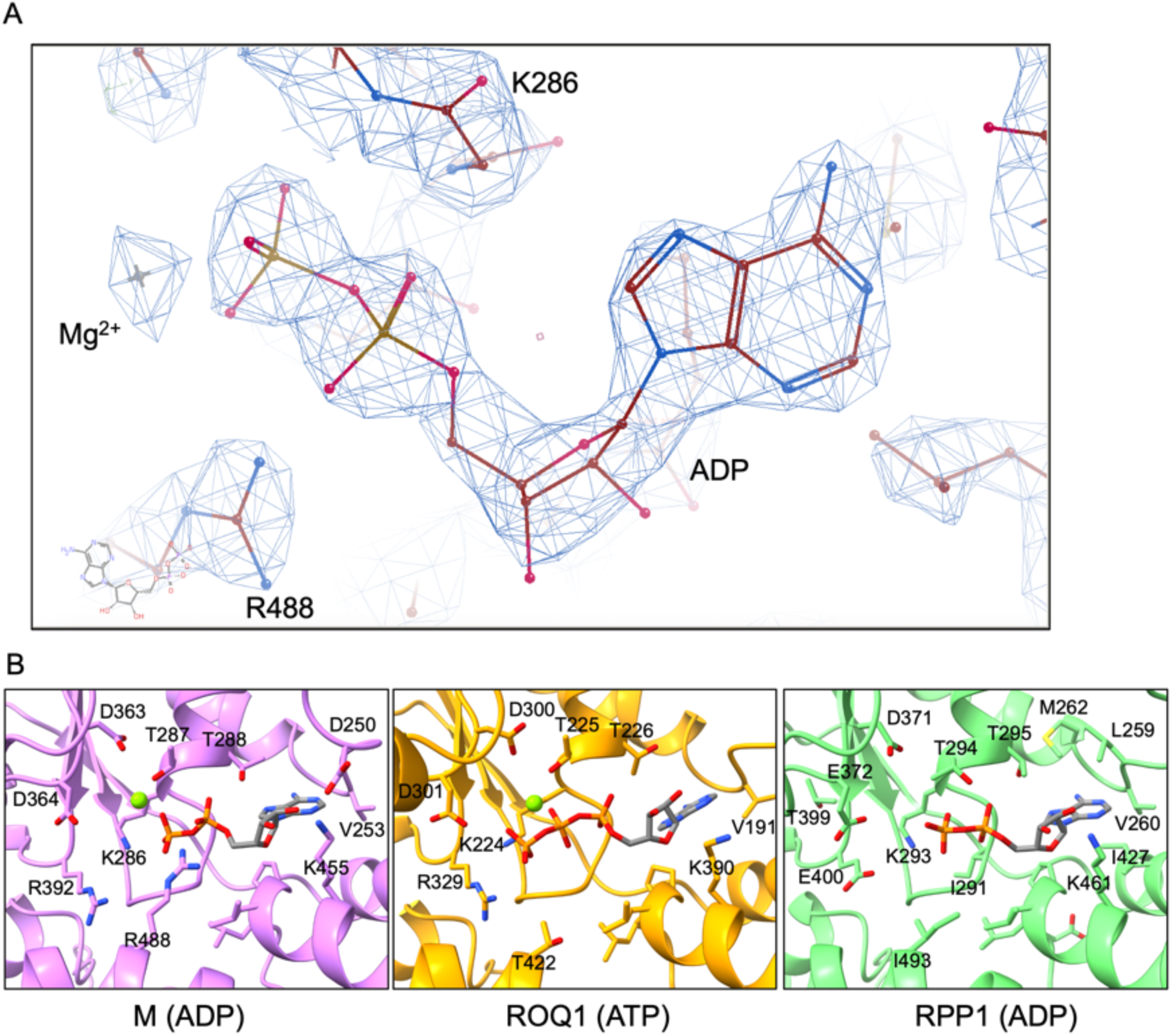
The ADP-bound state of the M resistosome. **(A)** Cryo-EM density of the ADP-bound NB-ARC region is shown with the model. No density was observed to account for the presence of an ATP gamma-phosphate moiety. **(B)** The nucleotide binding region of M (purple), ROQ1 (orange) and RPP1 (green) showing ADP, ATP and ADP binding, respectively.

**Figure S11.**
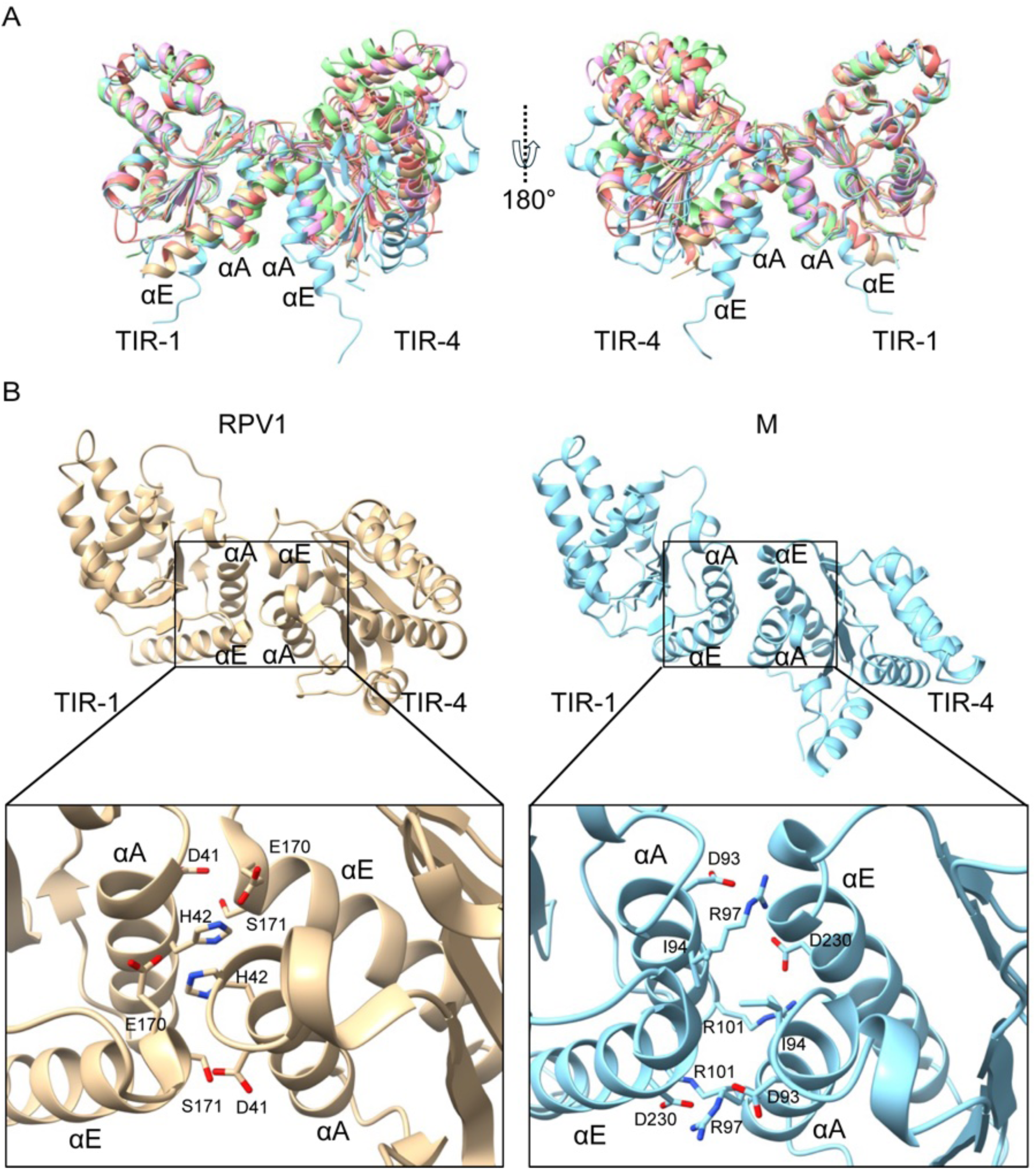
Structural analysis of the AE interface of TIR domains of the M resistosome. **(A)** Superposition of TIR symmetric AE interface-mediated homodimers from RUN1^TIR^ (salmon; PDB: 7S2Z), RPV1^TIR^ (brown; PDB: 5KU7), the ROQ1 resistosome (pink; PDB: 7JLX) and the RPP1 resistosome (green; PDB: 7CRC) onto the TIR symmetric homodimers of the M resistosome (cyan; PDB: 9PT6). The TIR-1 molecule was used for superposition. Only TIR-1 and TIR-4 molecules are shown. **(B)** Overview (top) and close-up view (bottom) of the AE interface from the RPV1 crystal structure (left; PDB: 5KU7) and the M resistosome (right; PDB: 9PT6).

**Figure S12.**
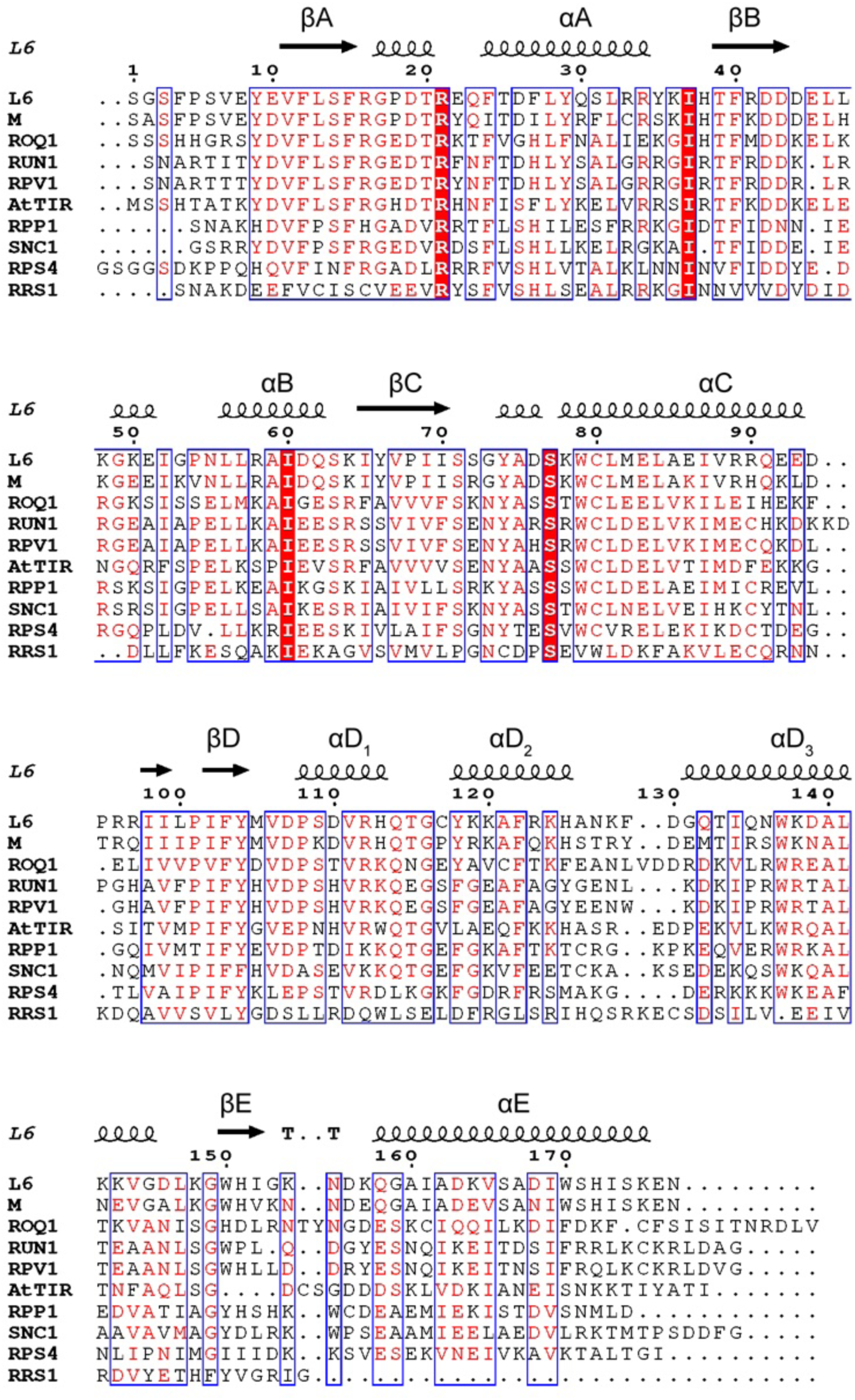
Multiple sequence alignment of plant TIR domains. The TIR domain from L6, M, ROQ1, RUN1, RPV1, AtTIR, RPP1, SNC1, RPS4 and RRS1 sequences were used for analysis. The alignment figure was generated using ESPript ^75^. The elements of secondary structure are shown above the alignment, based on the L6^TIR^ crystal structure (PDB: 3OZI).

**Figure S13.**
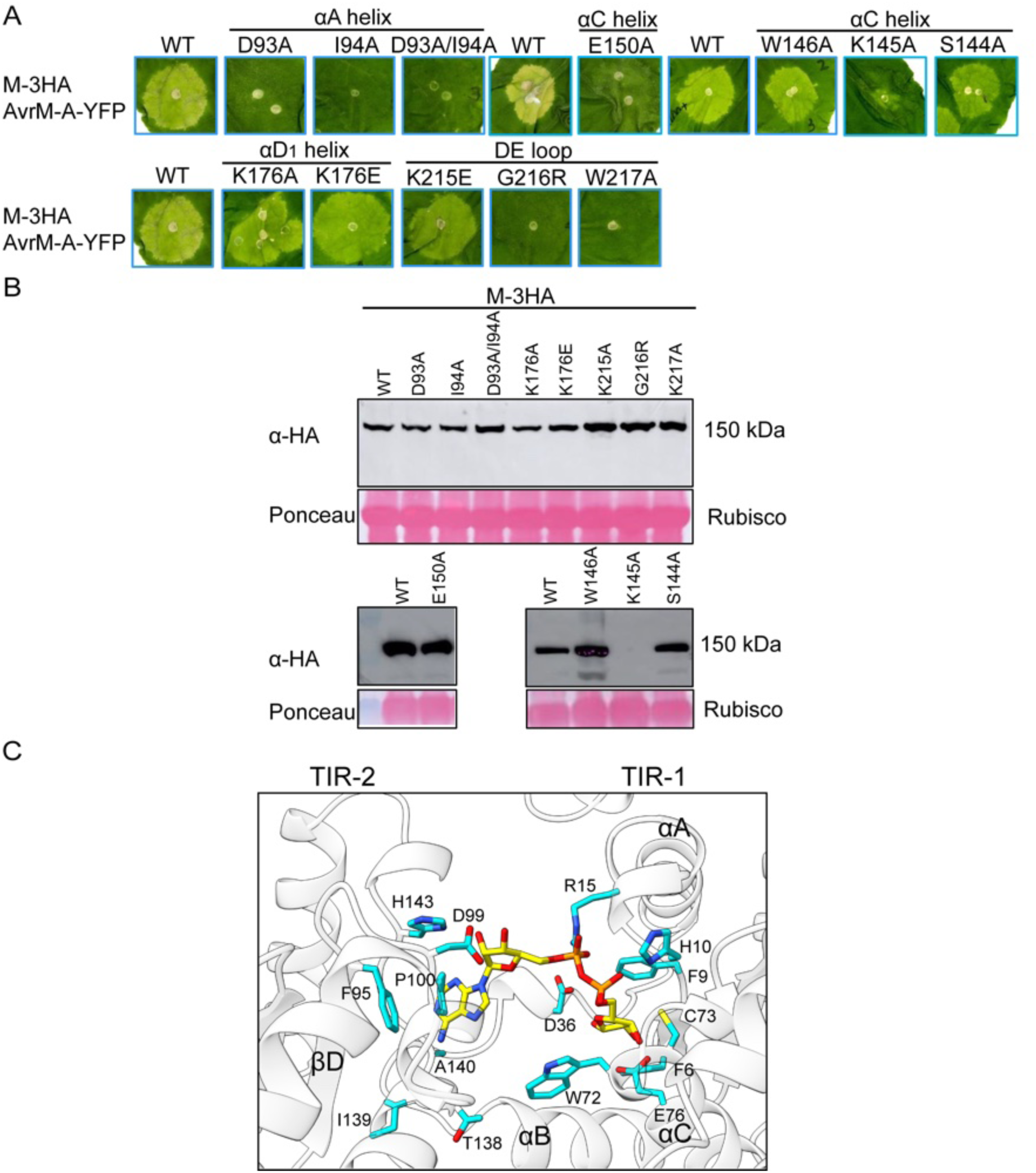
Substrate binding by plant TIR domains and effects of M mutations in the TIR domain on effector-dependent cell death. **(A)** Transient co-expression of AvrM-A (fused to yellow fluorescent protein (YFP)) with wild-type (WT) M (tagged with 3HA), or the M mutant constructs (D93A, I94A, D93A/I94A), by *Agrobacterium*-mediated infiltration of *N. benthamiana* leaves. **(B)** Immunoblot detection of M or mutant variants using anti-HA antibodies. Ponceau staining of the membrane, subsequently used for western blot, is also shown. **(C)** Crystal structure of RPP1^TIR^ domains bound to ADPR (PDB: 7XOZ) is shown in a close-up view. Conserved residues involved in substrate binding of RPP1^TIR^ are also observed for M^TIR^ and SARM1^TIR^ bound to 1AD in Figure 5D for comparison.

**Figure S14.**
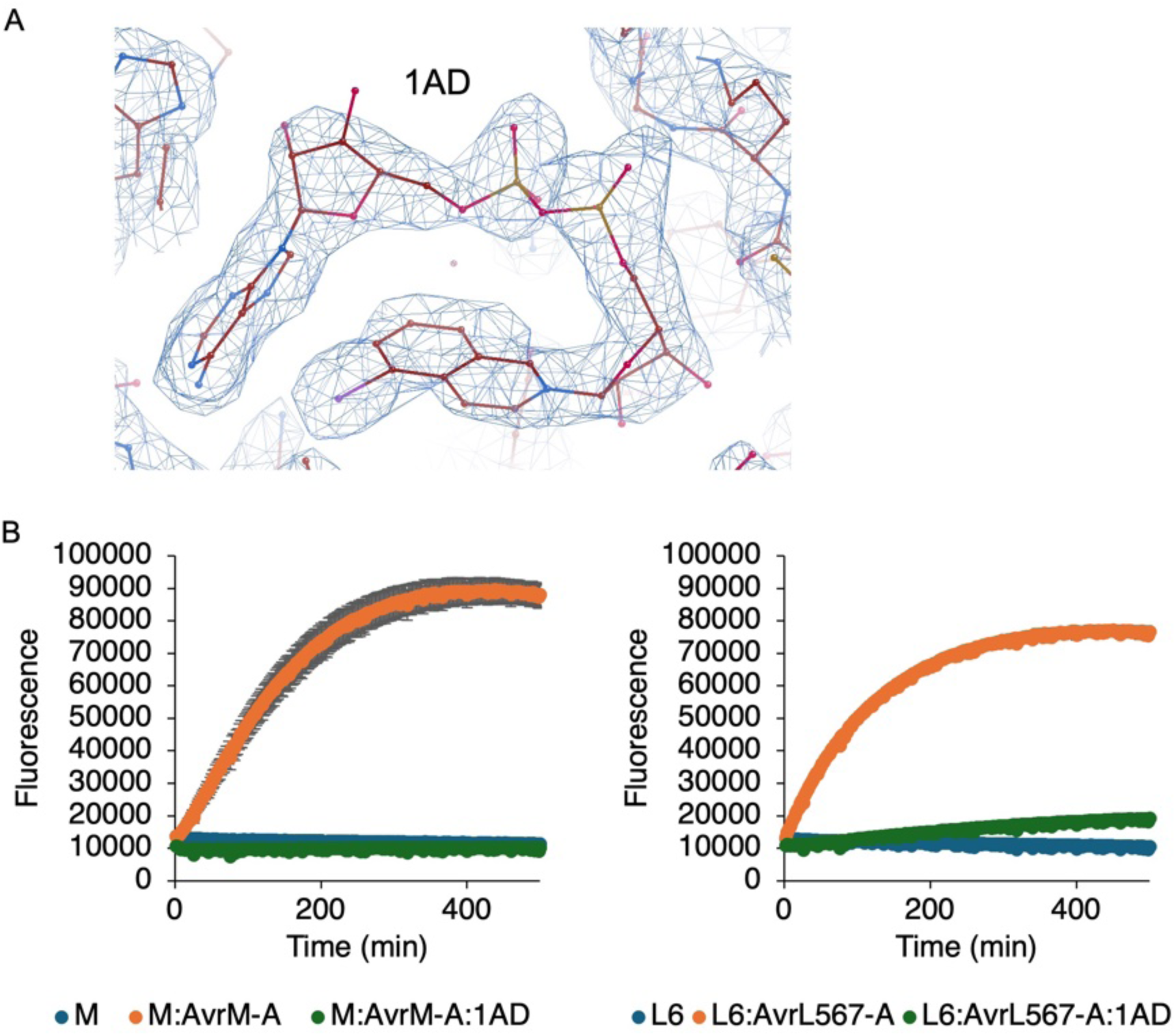
Fluorescence-based NADase assays for full-length flax M and L6 proteins and effects of 1AD on the NADase activity. **(A)** The atomic model and the cryo-EM density of the 1AD-binding region is shown. **(B)** Enzymatic activities of M (left) and L6 (right) full-length TNL proteins, cleaving 100 μM εNAD (fluorescent NAD^+^ analogue), were tested in the presence or absence of their cognate effectors (AvrM-A and AvrL567-A, respectively). 1 mM ATP was supplemented with reactions containing effectors. To test whether 1AD serves as an inhibitor of NADase activity, 400 μM 1AD was added to the reaction mixture and incubated at room temperature for 1.5 h, prior to adding εNAD.

**Figure S15.**
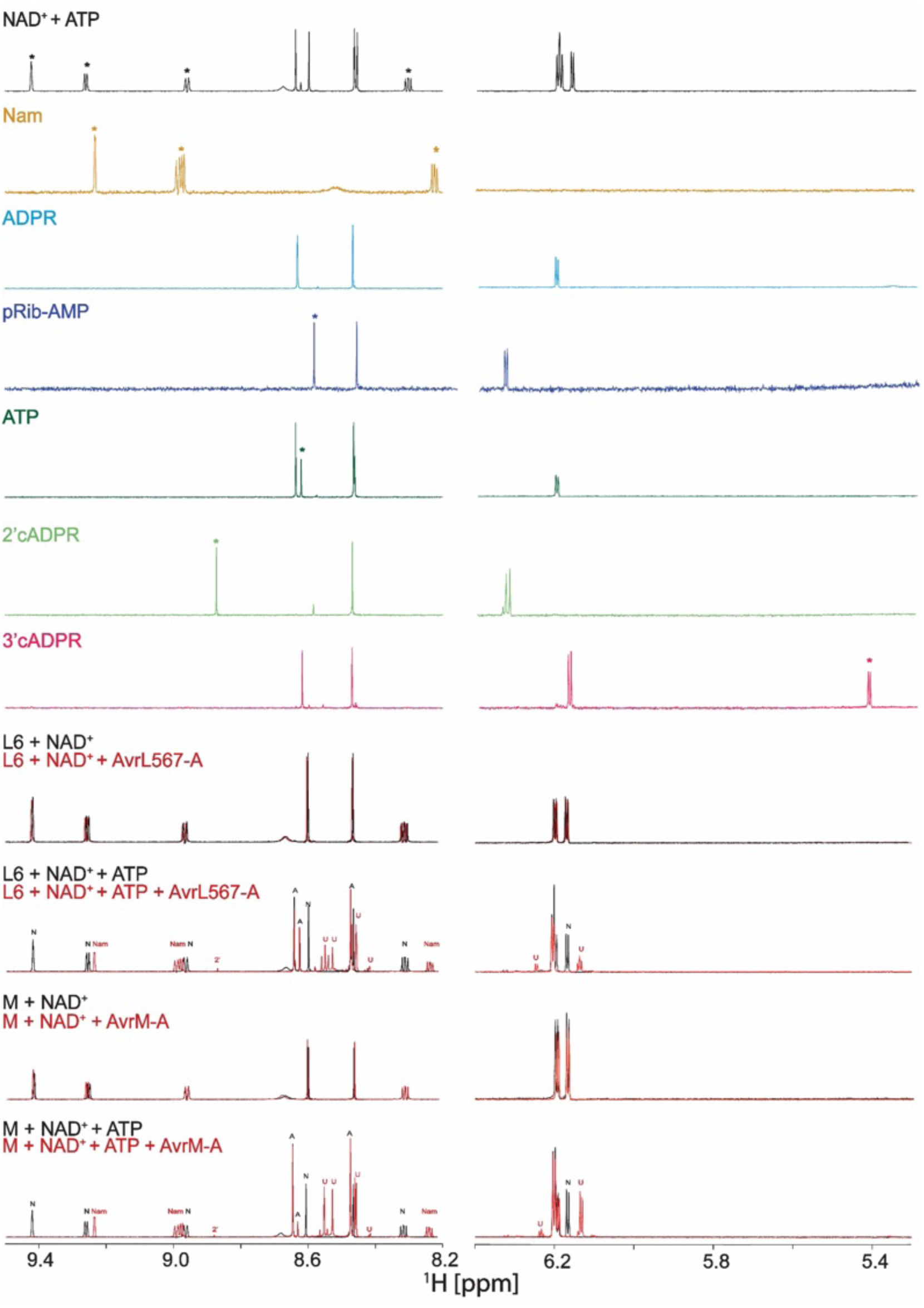
Extended ^1^H NMR spectral regions during NAD^+^ cleavage. The chemical shift region displaying the peaks of interest is shown (9.5 – 8.2 ppm and 6.4 – 5.3 ppm). Standards are shown, where asterisks indicate which peaks are unique and can be used to determine where a compound is present in test samples. M and L6 were incubated with different conditions, as described in the Methods section. Peaks are labeled with N (NAD^+^), Nam (nicotinamide), 2’ (2’cADPR), A (ATP) and U (unique/unknown – peaks that do not match any standard). Note that the peaks for ADPR and ATP overlap significantly and therefore we did not label any peaks with ADPR. pRib-AMP and 3’cADPR were not detected in any sample.

**Figure S16.**
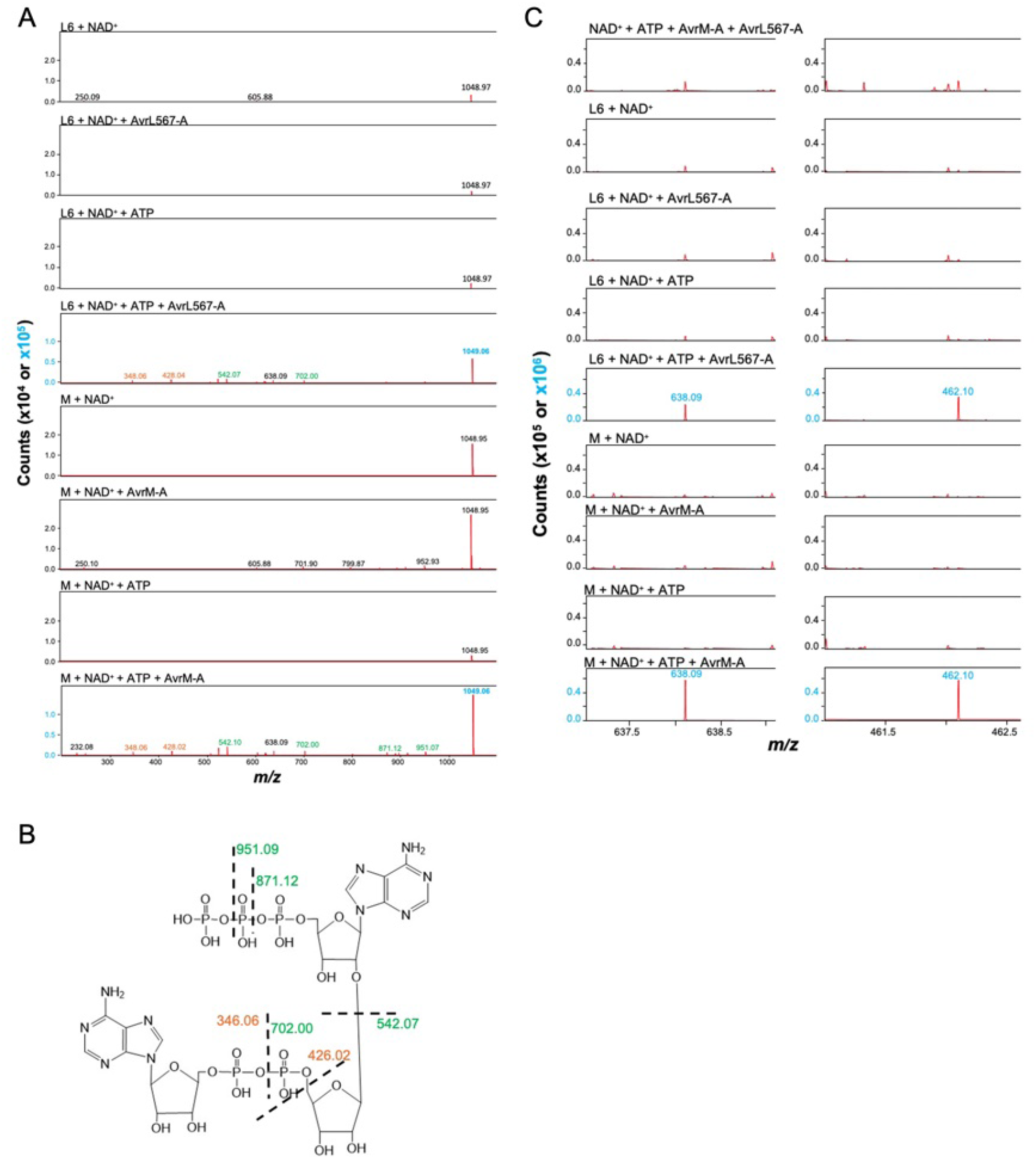
MS analysis of TNL enzyme assays. **(A)** MS/MS analysis using a range of 1049.1 +/- 0.4 *m/z*. Data for each sample were collected with a collision energy of 15 eV, to capture the parent compound, and data was summed with a second data collection, where collision energy was increased to 35 eV (to capture the fragments identified). The y-axis has a smaller range, to demonstrate a lack of ADPr-ATP and related fragments in samples lacking NADase activity. Only ions with an intensity of ≥ 5,000 counts were used. Black text indicates fragments not mapped. Green indicates ions that are positively charged upon fragmentation and gain no additional H. Yellow fragments indicated negatively charged fragments that gain 2H to become positively charged (M^-1^ + 2H^+^)^+^. **(B)** ADPr-ATP with fragments identified in panel (A). **(C)** Additional ions were detected in samples with enzyme activity, a 638.09 *m/z* (note this was a fragment observed in (A)) and a 462.1 *m/z* species.

**Figure S17.**
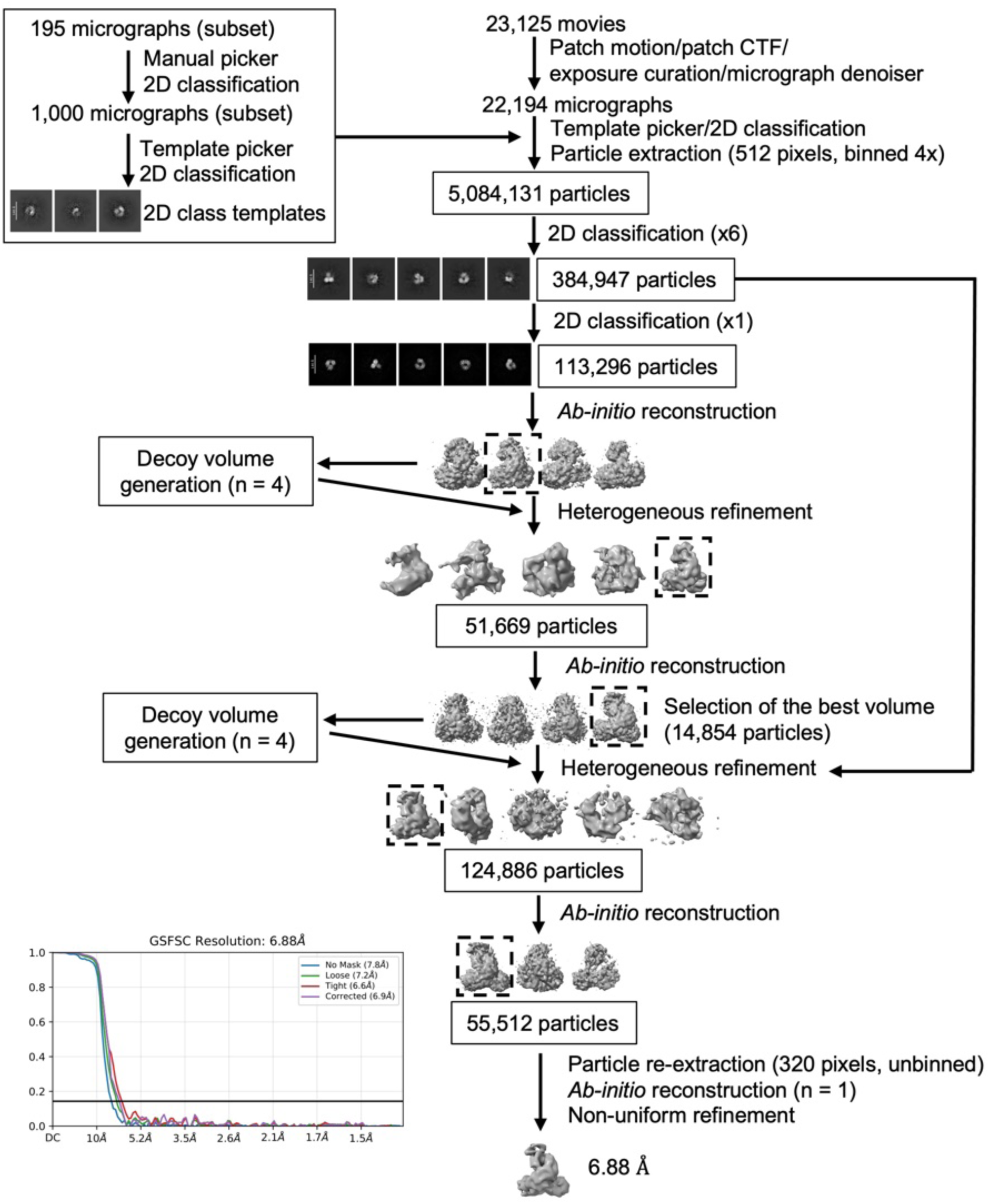
Cryo-EM SPA data processing workflow for the M monomer.

**Figure S18.**
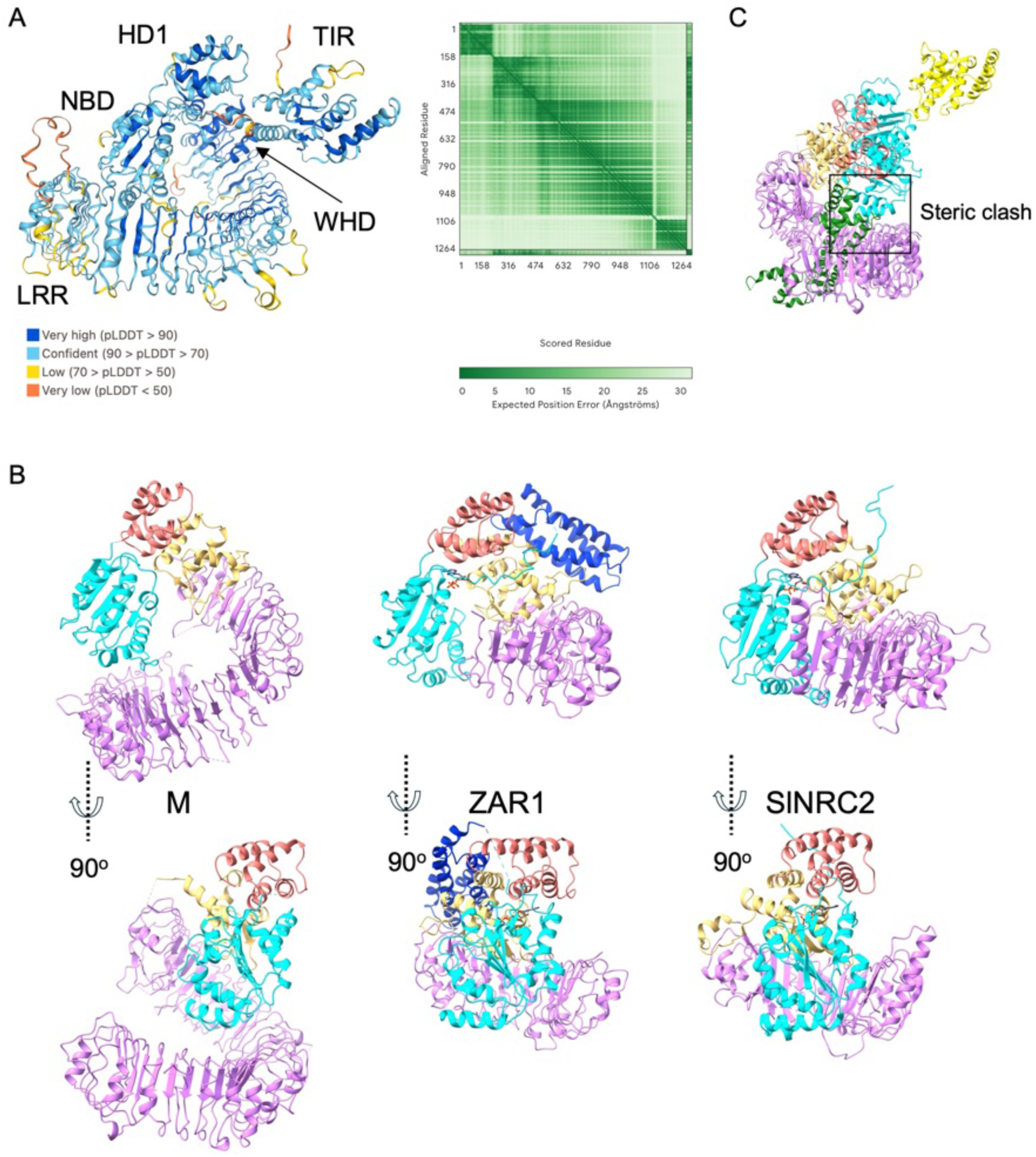
The M monomer protein structure. **(A)** AlphaFold 3-predicted structure of the M monomeric protein (residues 69-1305) in complex with ADP. The structure (left panel) is shown with colors representing pLDDT (predicted local distance difference test) scores, indicating high confidence for the overall model prediction. PAE (predicted aligned error) plot is shown on the right panel; ipTM = 0.97; pTM = 0.61. **(B)** Structure comparison between the cryo-EM structure of the M monomer, ZAR1 monomer (PDB: 6J5W), and the protomer of dimeric *S. lycopersicum* NRC2 (PDB: 8XUO), shown side-by-side at two different angles (90° rotation in the bottom panel). ADP is bound to ZAR1 and NRC2. The CC domain of the ZAR1 monomer is resolved, while that of NRC2 is undefined, likely due to flexibility of this region. **(C)** Superposition of the M monomer onto the protomer of M bound to AvrM-A from the M resistosome shows steric clashes between the NBD and AvrM-A.

**Figure S19.**
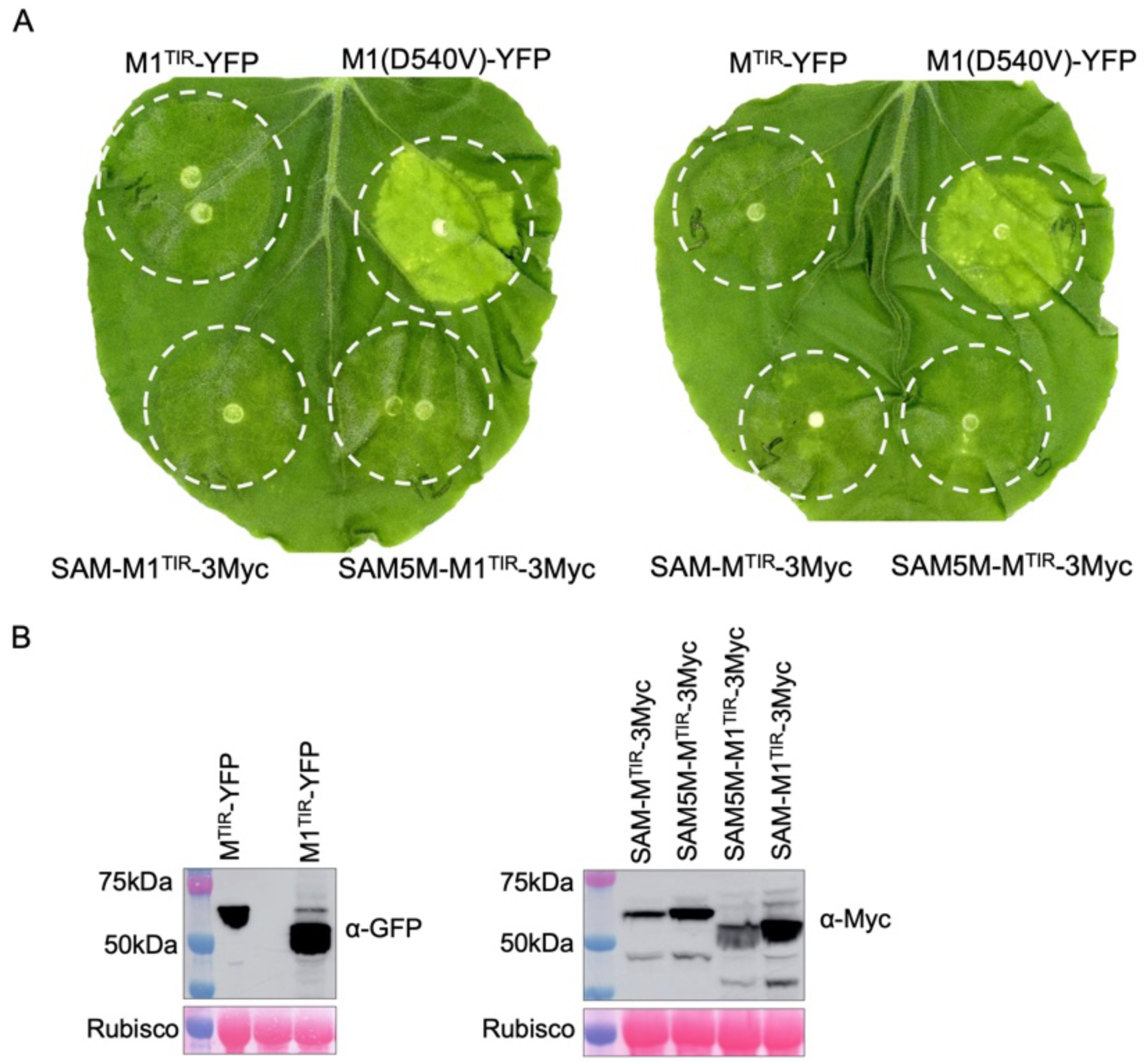
TIR domains of flax M and M1 do not trigger cell death when transiently expressed in leaves of *N. benthamiana*. **(A)** Visual evaluation of cell death upon transient expression. Constructs tested include: the N-terminal region of M (residues 1-263) and the related M1 (residues 1-248) that were C-terminally fused to YFP (yellow fluorescent protein) or N-terminally fused to hSARM1^tSAM^ or hSARM1^tSAM(5Mut)^ with a C-terminal Myc-tag; an autoactive full-length M1 mutant (D540V) C-terminally fused to YFP was used as a positive control. **(B)** Immunoblot analysis indicates that the M and M1 TIR domains are expressed when fused to YFP (α-GFP antibodies) or SAM domains (α-Myc antibodies).

**Table S1.**
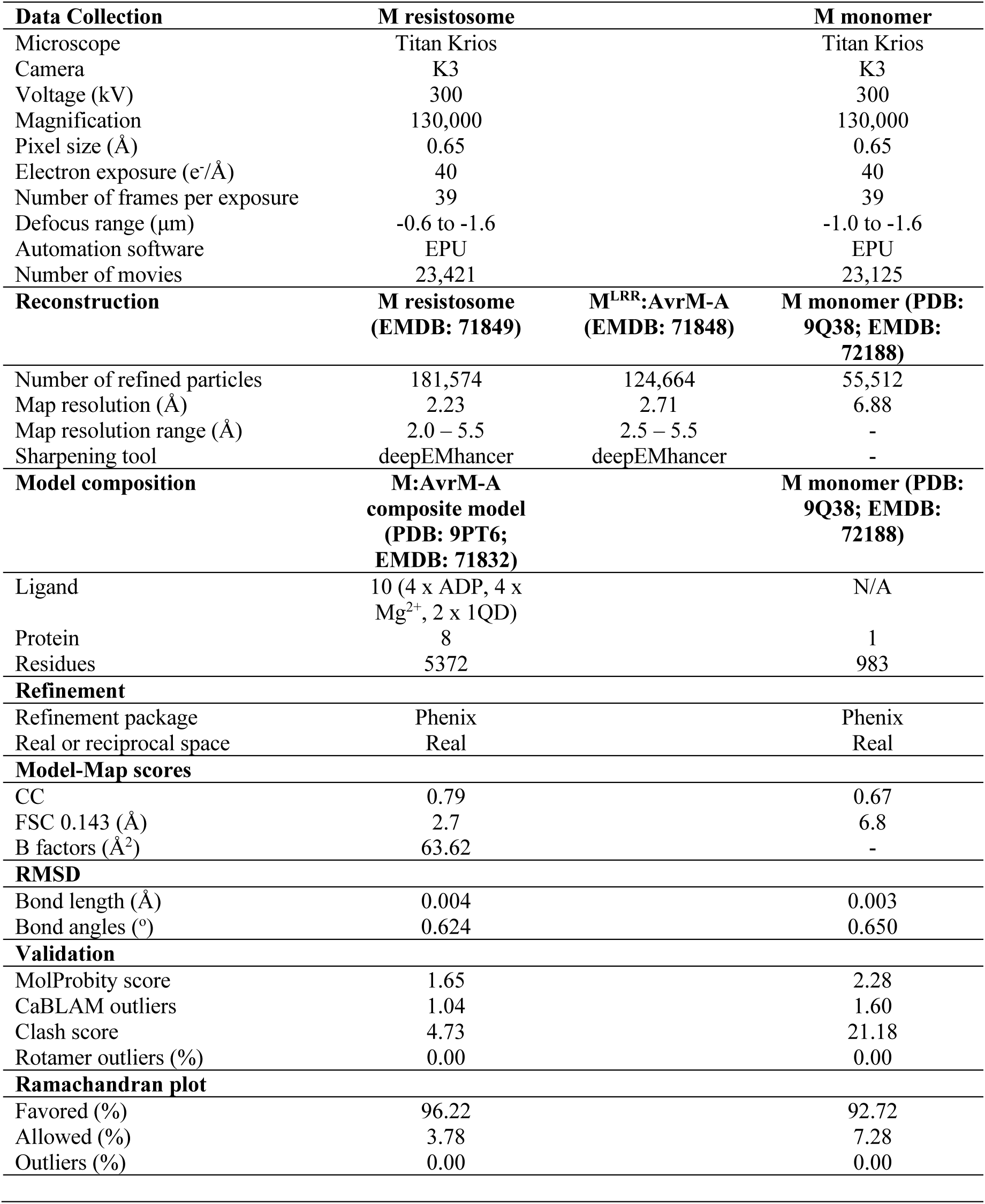
Statistics of the cryo-EM data collection, data processing and model refinement.

